# Antigen-specific induction of CD4^+^CD8αα^+^ intraepithelial T lymphocytes by *Bacteroidetes* species

**DOI:** 10.1101/2020.08.05.236513

**Authors:** Djenet Bousbaine, Preksha Bhagchandani, Mariya London, Mark Mimee, Scott Olesen, Mathilde Poyet, Ross W. Cheloha, John Sidney, Jingjing Ling, Aaron Gupta, Timothy K. Lu, Alessandro Sette, Eric J. Alm, Daniel Mucida, Angelina M. Bilate, Hidde L. Ploegh

## Abstract

The microbiome contributes to the development and maturation of the immune system^1–3^ In response to commensal bacteria, CD4^+^ T cells can differentiate into distinct functional subtypes with regulatory or effector functions along the intestine. Peripherally-induced Foxp3^+^-regulatory T cells (pTregs) maintain immune homeostasis at the intestinal mucosa by regulating effector T cell responses against dietary antigens and microbes^4^. Similarly to pTregs, a subset of small intestine intraepithelial lymphocytes CD4^+^CD8αα^+^ (CD4_IELS_) exhibit regulatory properties and promote tolerance against dietary antigens^5^. Development of CD4_IELS_ from conventional CD4^+^ T cells or from Treg precursors depends on the microbiota^5,6^. However, the identity of the microbial antigens recognized by CD4_IELs_ remains unknown. We identified species belonging to the *Bacteroidetes* phylum as commensal bacteria capable of generating CD4_IEL_ from naïve CD4^+^ T cells expressing the pTreg transnuclear (TN) monoclonal TCR^6^ as well as from polyclonal WT T cells. We found that β-hexosaminidase, a widely conserved carbohydrate-metabolizing enzyme in the *Bacteroidetes* phylum, is recognized by TN T cells, which share their TCR specificity with CD4^+^ T cells found in the intraepithelial compartment of polyclonal specific-pathogen-free (SPF) mice. In a mouse model of colitis, β-hexosaminidase-specific CD4_IELs_ provided protection from ulceration of the colon and weight loss. Thus, a single T cell clone can recognize a variety of abundant commensal bacteria and elicit a regulatory immune response at the intestinal epithelial surface.

## Main text

The microbiota contributes to functional specification of adaptive immunity, both through direct interactions and via soluble mediators released into the environment. In turn, adaptive and innate immunity shape the microbiota, for example through production of antibacterial peptides and antibodies. The protective functions of T cells induced by microbes range from antibacterial defense to cancer immunity^7^ and assistance in wound healing^8^, but diseases such as colitis can ensue when these interactions are perturbed. The microbiota shapes the plasticity and adaptation of CD4^+^ T cells in the intestinal lamina propria. Colonic bacteria such as *Helicobacter hepaticus* promote differentiation of antigen-specific CD4^+^ T cells into Foxp3^+^ regulatory T cells (Tregs) in the colon, while Segmented Filamentous Bacteria (SFB) induce quasi-clonal pro-inflammatory TH17 cells in the ileum^9–13^. These studies highlight the importance of a specialized and diverse intestinal immune response to commensals and pathobionts localized in different intestinal niches. Such interactions are not only species-specific, but depending on the anatomical sites where they occur, can influence T cell fates^6^. Ablation of SFB eliminates the corresponding TH17 T cells^14^, while Class Ib MHC-restricted CD8^+^ T cells require the presence of *S. epidermidis* in the skin^15^. Fate decisions must thus be made at the clonal level and different T-cell receptor (TCR) specificities ought to drive distinct developmental and functional outcomes.

Two important subsets of CD4^+^ lymphocytes regulate adaptive immunity at the intestinal mucosa: peripheral regulatory T cells (pTregs) and double-positive (CD4^+^CD8αα^+^) intraepithelial lymphocytes (IEL), hereafter referred to as CD4_IELs_. Maintenance of immune homeostasis at the intestinal mucosa requires the action of pTregs^16,17^, which regulate the response of effector T cells against dietary antigens and commensals. Small intestine CD4_IELs_ likewise promote tolerance to dietary antigens^5,18^. Depletion of CD4_IELs_ causes disease in mice that lack functional Foxp3^5^. CD4_IELs_ and pTregs thus cooperate in the regulation of local intestinal inflammation. In specific-pathogen-free (SPF) mice, CD4_IELs_ are present mostly in the epithelium of the small intestine. Their abundance varies with both age and diet and depends on the indigenous microbiota^19–21^ (Extended Data Fig. 1a,b). CD4_IELs_ can develop either from conventional CD4^+^ T cells or from Foxp3^+^ Treg precursors in a microbiota-dependent manner^5,6^. Germ-free (GF) mice have few if any CD4_IELs_, regardless of their genetic background^5,22^ (Extended Data Fig. 1c). Mice treated with antibiotics^5,22^ show a similar decline in CD4_IELs_ (Extended Data Fig. 1d). Fecal transplants from mice housed in different SPF facilities restore the CD4_IEL_ compartment in GF mice to different extents^19,20^ (Extended Data Fig. 1e). Secretion of metabolites by certain commensals can further promote accumulation of CD4_IELs_^19^. The microbiota therefore contributes not only to the development but also to the maintenance of CD4_IELs_. However, the identity(ies) of the microbial antigens recognized by CD4_IEL_ TCRs is unknown. Most IEL populations, including CD4_IELs_, appear to have a somewhat restricted TCR repertoire^19^,^23–29^. A limited diversity of antigens might thus suffice to shape this compartment, raising the possibility that a common antigen is recognized by CD4_IELs_ TCRs. We used the microbiota-specific transnuclear (TN) monoclonal model^6^ to identify naturally-occurring TCR ligands of CD4_IELs_. T cells from the TN mouse carry a monoclonal TCR cloned from a pTreg derived from the intestine-draining mesenteric lymph nodes (mLN)^13^. Upon transfer, naïve T cells isolated from TN mice populate not only the recipient’s mLNs, where they can differentiate into pTregs, but also the epithelium of the small intestine, where they convert into CD4_IELs_^6^ (Extended Data Fig. 2a,b). TN CD4^+^ T cells require the presence of the microbiota to reside, persist and differentiate into CD4_IELs_ at that location^6^.

We thus sought to identify the commensal member(s) recognized by the TN cells that allow them to migrate and convert into CD4_IELs_ at the intestinal epithelium. The microbiota of mice sourced from Taconic was enriched for the antigen recognized by the TN TCR^6^. To identify the species recognized by the TN TCR, we grew fecal bacteria derived from Taconic excluded flora (EF) mice under various culture conditions to select for a diverse array of culturable bacteria. We then co-cultured TN T cells with dendritic cells (DCs) in the presence of sonicated bacterial extracts obtained from different bacterial cultures. We observed strong proliferation of TN T cells in the presence of bacterial extracts isolated from Bacteroides bile esculin (BBE) agar plates or aerotolerant bacteria isolated from Schaedler blood agar plates (Fig. 1a,b). 16S rRNA sequencing showed enrichment for operational taxonomic units (OTUs) of the *Parabacteroides* genus in the cultures that yielded proliferation of TN cells (Fig. 1c). We tested several *Parabacteroides* strains isolated from mice and humans and observed extensive proliferation of TN T cells in the presence of *Parabacteroides goldsteinii* extracts, the predominant member of the Altered Schaedler Flora (ASF)^30^ (Fig. 1d and Extended Data Fig. 2c). Proliferation of TN T cells in response to *P. goldsteinii* protein-rich extract was Class II MHC-restricted and dependent on antigen presentation by dendritic cells^6^ (Extended Data Fig. 2d). Neither the related *Parabacteroides distasonis* nor any other species tested induced proliferation of TN T cells *in vitro* (Fig. 1d, Extended Data Fig. 2e,f). To determine whether *P. goldsteinii* extracts could also induce TN T cell proliferation *in vivo,* we transferred CD4^+^ TN T cells into congenically-marked recipients that were then immunized subcutaneously with bacterial extracts in alum. Robust proliferation of TN T cells occurred in the draining lymph nodes (inguinal lymph nodes; iLNs) of mice that received *P. goldsteinii* extract but not *P. distasonis* extract or PBS (Fig. 1e,f). thus unequivocally identifying *P. goldsteinii* as a bacterium capable of engaging the pTreg TN TCR.

**Figure 1:**
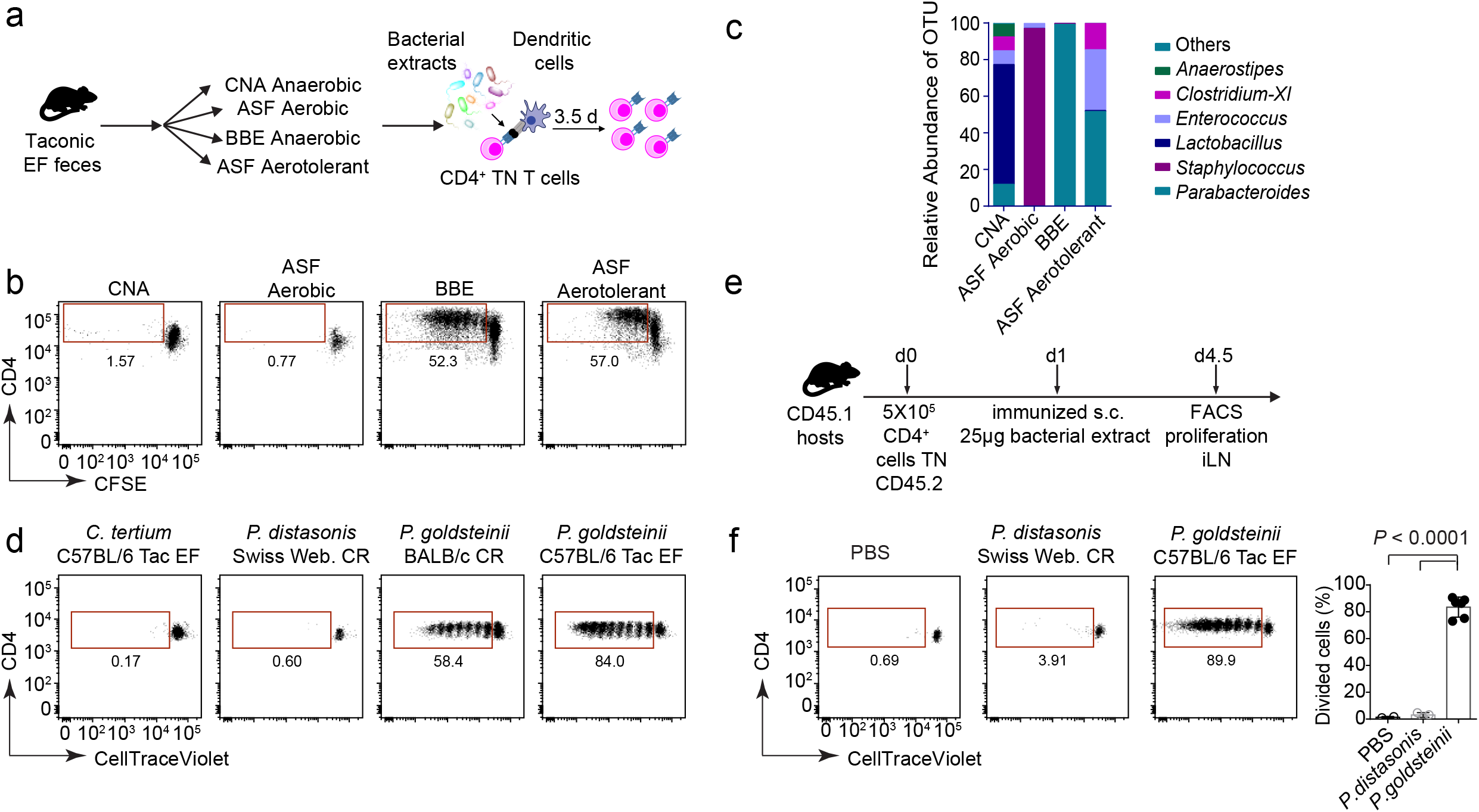
The pTreg TN TCR recognizes *P. goldsteinii*. (**a**) Schematic of the *in vitro* proliferation assay. (**b**) Naïve CD4^+^ TN T cells were labeled with CFSE and co-cultured with dendritic cells (DCs) purified from B16-Flt3L-injected mice in the presence of bacterial extracts derived from Taconic EF fecal bacteria, isolated using the indicated growth conditions. After 3.5 days, CFSE dilution was assessed by flow cytometry. Dot plots are representative of at least 3 experiments. Cells were gated on Via-probe^-^ CD4^+^ Vα2^+^ Vβ6^+^. (**c**) DNA from the bacterial extracts depicted in (a) was isolated, the V4 region of 16s rRNA was amplified using nested PCRs and sequenced using Illumina technology. (**d**) *In vitro* proliferation as in (a): co-culture of CellTrace Violet-labeled naïve CD4^+^ TN T cells and DCs purified from B16-Flt3L-injected mice in the presence of the indicated bacterial extracts. Dot plots are representative of at least 3 experiments. (**e**) Schematic of the *In vivo* proliferation assay shown in (f). Congenically marked WT CD45.1^+^ mice received 7X10^5^ CellTrace-Violet-labeled naïve CD45.2^+^ CD4^+^ TN T cells and were immunized subcutaneously with 25μg of bacterial extracts adsorbed in alum 1 day later. 3.5 days after immunization, proliferation was assessed in the draining lymph node (inguinal lymph node, iLN) by flow cytometry. (**f**) Dot plot shows dilution of CellTrace-Violet in CD45.1^-^ CD45.2^+^CD4^+^Vα2^+^Vβ6^+^ T cells in one representative mouse per condition. The graph shows all mice analyzed in two independent experiments (PBS n=2, *P. distasonis* n=3, *P. goldsteinii* n=6). The graph shows means +/− standard deviation and each symbol represents a single mouse. P values were calculated using unpaired one-sided Student’s t tests. P values <0.05 were considered statistically significant. Tac=Taconic, CR=Charles River, Web.=Webster.

We next asked whether colonization with *P. goldsteinii* could induce proliferation of naïve CD4^+^ TN T cells and their conversion into CD4_IELs_ *in vivo.* We adoptively transferred TN T cells into RAG-deficient recipients pre-treated with antibiotics^31^, which we then colonized with *P. goldsteinii* (Fig. 2a). Colonization was confirmed by *P. goldsteinii-* specific qPCR (Extended Data Fig. 3a). Mice treated only with antibiotics or colonized with unrelated bacteria *(Clostridium tertium)* isolated from the cecum of mice that served as the source of *P. goldsteinii,* failed to expand and convert TN cells (Fig. 2b-d). In contrast, in mice colonized with *P. goldsteinii,* TN cells proliferated extensively and converted into CD4_IELs_ (Fig. 2b-d). We conclude that *P. goldsteinii* promotes the development of CD4_IELs_ from TN T cells *in vivo.*

**Figure 2:**
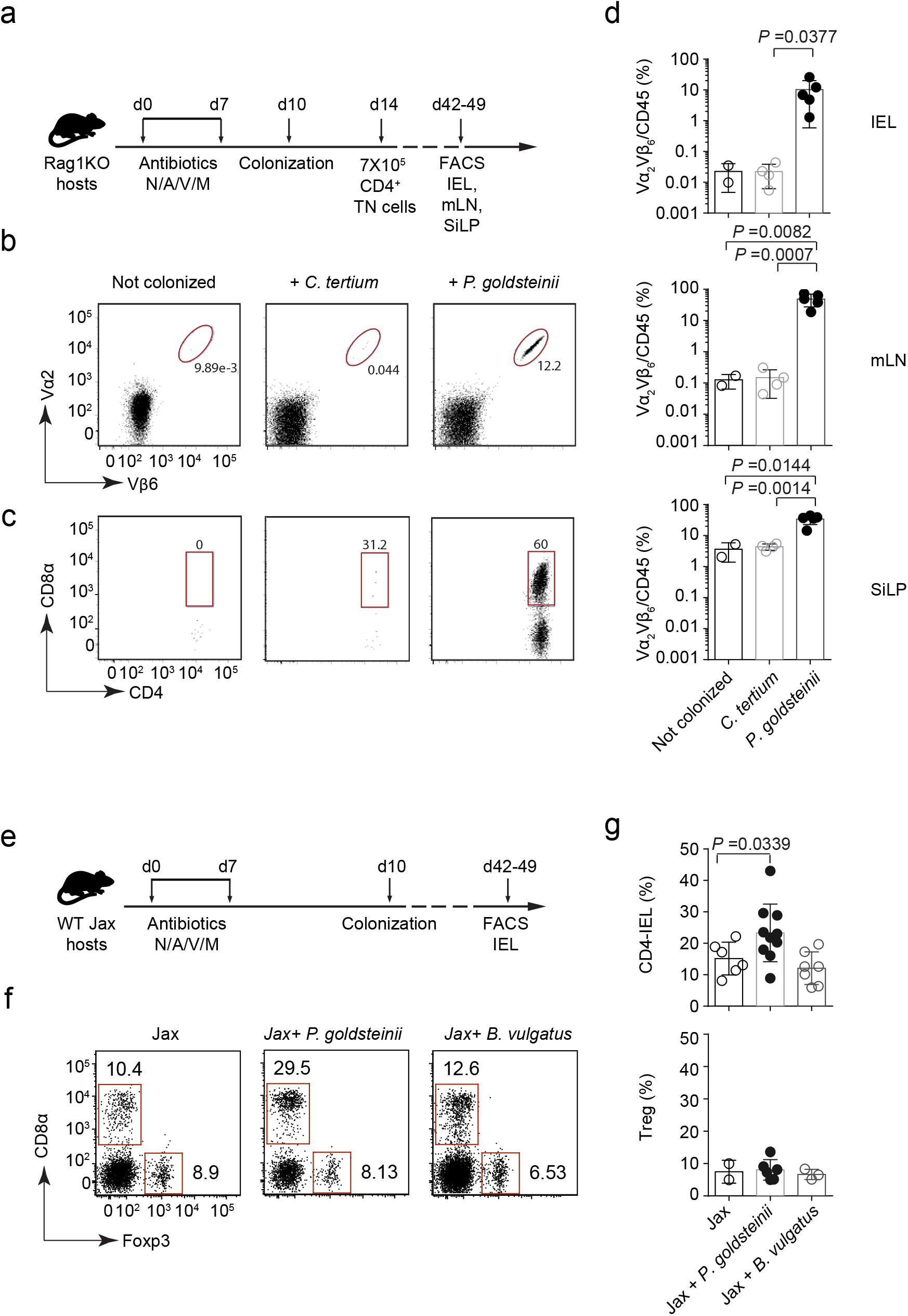
*P. goldsteinii* induces CD4_IELs_ in both monoclonal and polyclonal SPF mice. (**a**) Experimental design. Rag1KO hosts were treated with antibiotics (ABX) for 7 days. At day 10, recipient mice were colonized or not (n=2) with the indicated bacteria *(C. tertium* n=4, *P. goldsteinii* n=5). The following day, mice received 7X10^5^ naïve CD45.2^+^ CD4^+^ TN T cells. On day 42-49, the small intestine intraepithelial compartment (IEL), small intestine lamina propria (SiLP) and the mesenteric lymph nodes (mLNs) were harvested. We analyzed their cellular composition by flow cytometry. (**b**) Dot plots show the frequency of TN cells (Vα2^+^Vβ6^+^) among CD45^+^ cells of one representative mouse per group. (**c**) Dot plots shows the frequency of CD4_IELs_ among TN cells in one representative mouse per group. Cells were gated on CD45^+^ CD4^+^ Vα2^+^Vβ6^+^. (**d**) Graphs show all mice analyzed in two independent experiments shown in (b,c). The graphs show the mean +/− standard deviation (SD) and each symbol represents a single mouse. P values were calculated using unpaired one-sided Student’s t tests. P values <0.05 were considered statistically significant. (**e**) Experimental design. WT Jax mice were treated with ABX for 1 week. Three days later they were colonized with the indicated bacteria (Jax n=6, *P. golsteinii* n=10, *B. vulgatus* n=7). At day 42-49, IELs were analyzed by flow cytometry. (**f**) Dot plots show the frequency of WT cells in the IEL of one representative mouse per group. Cells were gated on live Aqua ^-^CD45^+^TCRγδ^-^ TCRβ^+^CD8β^-/lo^CD4^+^. (**g**) Graphs show all mice analyzed in two independent experiments performed in two different animal facilities shown in (e). The frequency of both CD4_IELs_ and Tregs was assessed in the small intestine intraepithelial compartment. The graphs show the means +/− SD and each symbol represents a single mouse. P values were calculated using unpaired onesided Student’s t tests. P values <0.05 were considered statistically significant.

*P. goldsteinii* is prevalent in many animal facilities^32^, but less so in mice from Jackson Laboratories (Jax) (Extended Data Fig. 3b,c) which also harbor fewer CD4_IELs_ than mice from other facilities (Extended Data Fig. 1a). Indeed, we could not detect any *P. goldsteinii* 16s read in the feces of Jax mice in contrast to mice housed at RU (Supplementary Data Table 1). To test whether the CD4_IEL_ compartment of WT Jax mice could be boosted by *P. goldsteinii,* we colonized them with *P. goldsteinii* or with the related species *B. vulgatus,* or with the unrelated SFB (Fig. 2e-g and Extended Data Fig. 3d,e). While CD4_IELs_ were increased in mice colonized with *P. goldsteinii,* mice colonized with *B. vulgatus* or SFB showed similar frequencies of CD4_IELs_ as non-colonized mice. Thus, *P. goldsteinii* promotes the accumulation of CD4_IELs_ not only in the pTreg TN monoclonal model but also in WT polyclonal mice. However, in contrast to supplementation with *P. goldsteinii* in antibiotic-treated mice, colonization of fully germ-free (GF) mice with *P. goldsteinii* alone failed to expand CD4_IELs_ (Extended Data Fig. 3f). Thus, although recognition of *P. goldsteinii* antigens can promote CD4_IEL_ differentiation from naïve precursors, other commensals are required for full differentiation into the CD4_IEL_ phenotype in GF mice.

To identify the TCR ligands produced by *P. goldsteinii* that can induce TN and WT cells to become CD4_IELs_, we undertook a biochemical approach. We fractionated *P. goldsteinii* lysate by ammonium sulfate precipitation, followed by anion and cation exchange chromatography (Fig. 3a-d). These separations yielded fractions that were highly stimulatory to TN cells as assessed by *in vitro* proliferation assays (Fig. 3b-d and Extended Data Fig. 4a). Analysis of the most highly stimulatory fraction by LC-MS/MS identified 33 proteins with excellent tryptic peptide coverage (Extended Data Fig. 4b). The top 15 candidates from this list were expressed recombinantly in *Escherichia coli,* and sonicates prepared from the resulting *E. coli* recombinants served as a source of candidate antigens in the antigen presentation assay (Fig. 3a,e,f). This strategy identified β-N-acetylhexosaminidase (β-hex) as the protein recognized by the TN TCR. The β-hex gene is part of a polysaccharide utilization locus (Extended Data Fig. 4c), suggesting its involvement in the digestion of complex glycoproteins, either host or diet-derived^33^. Homologous sequences of this gene can be found across several bacterial phyla (Supplementary Data Table 2). TN T cells proliferated extensively in the draining lymph nodes of recipient mice immunized with a protein extract of *E. coli* expressing *P. goldsteinii* β-hex, but not when immunized with β-hex derived from closely related species (Fig 3g,i). Truncation analysis defined a ~70 residue stretch that contained the cognate epitope of the pTreg TN TCR (Extended Data Fig. 5a). Overlapping peptides of this region identified the epitope in a 14 amino acid stretch (Extended Data Fig. 5b). Taking advantage of the knowledge of anchor residues for I-A^b34^, we identified the minimal epitope as YKGSRVWLN (Extended Data Fig. 5c,d). We confirmed YKGSRVWLN as the cognate epitope of TN T cells by the *in vitro* and *in vivo* proliferative response of TN T cells to its synthetic version, both by sub-cutaneous and intranasal injections (Fig. 3h,i and Extended Data Fig 5c,e,f). Homologous synthetic β-hex peptides from *Parabacteroides merdae* failed to stimulate TN T cells. Lack of proliferation of TN T cells in response to the 14-mer N-acetylated *P. merdae* peptide was not due to an inability to bind class II MHC, as it was shown to bind I-A^b^ with intermediate affinity, albeit at about a 3-fold lower level than the *P. goldsteinii* epitope (Extended Data Fig. 5g).

**Figure 3:**
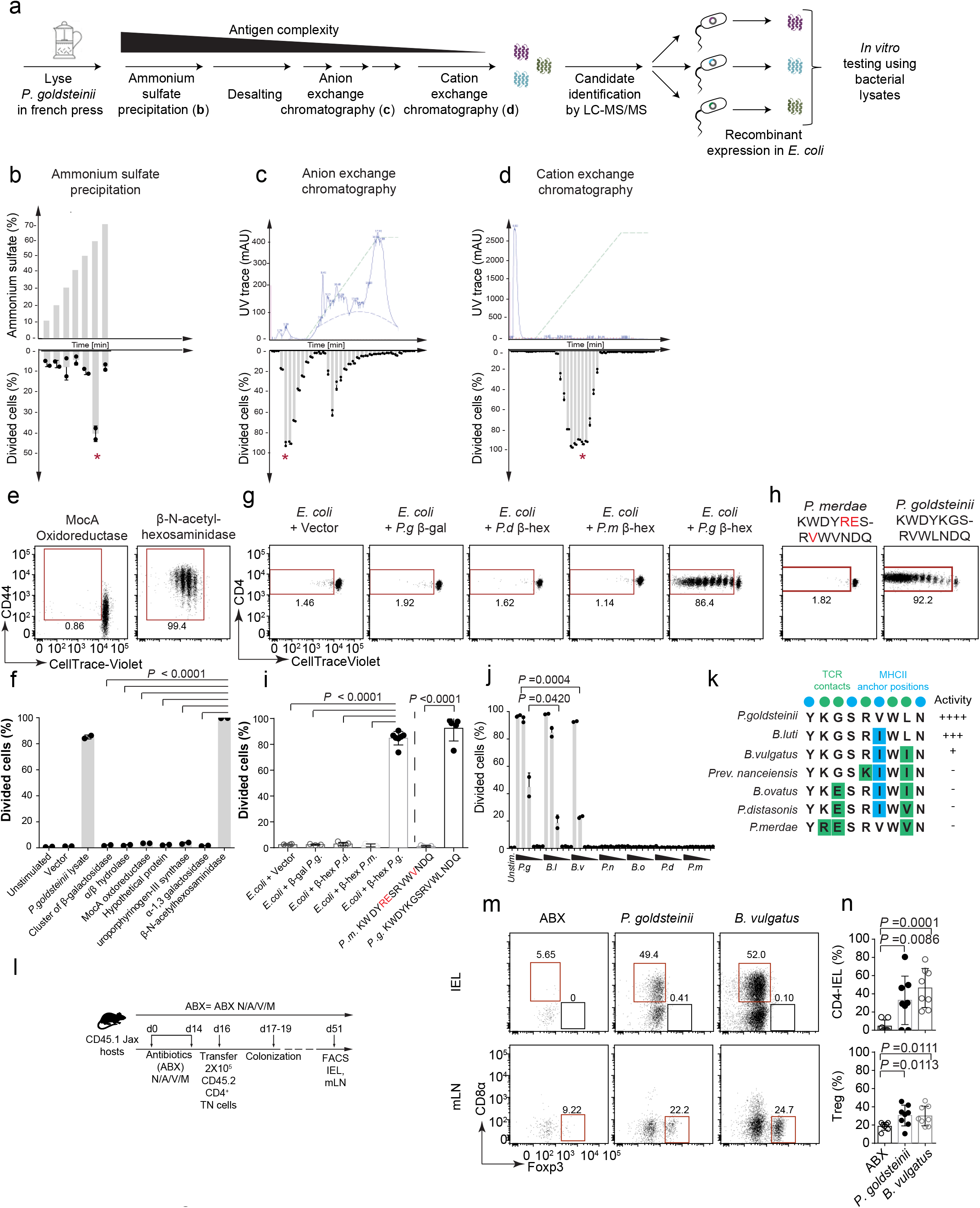
The TN TCR specifically recognizes epitopes from *Bacteroidetes* β-N-acetylhexosaminidase in complex with I-A^b^. (**a**) Experimental design. *P. goldsteinii* was grown anaerobically and lysed using a French press. Antigen recognized by the TN TCR was isolated by fractionating the lysate using a combination of ammonium sulfate precipitation (b), anion (c) and cation (d) exchange chromatography. Candidate polypeptides in the strongly stimulatory fractions were identified by mass spectrometry. The candidate polypeptides with the greatest sequence coverage and highest spectral counts were then recombinantly expressed in *E. coli* and tested individually *in vitro* for their ability to induce proliferation of TN cells (e,f). (**b**) *P. goldsteinii* lysate was fractionated using ammonium sulfate precipitation. Each fraction was desalted and tested for the presence of the antigen *in vitro:* naïve CD4^+^ TN T cells were labeled with CellTrace-Violet and co-cultured with dendritic cells purified from B16-Flt3L-injected mice in the presence of the indicated fractions. 3.5 days later, CellTrace-Violet dilution was assessed by flow cytometry. The graph shows the mean and standard deviation (SD) of one representative of at least two independent experiments. Each symbol represents one technical replicate. Cells were gated on Via-probe^-^CD4^+^Vα2^+^Vβ6^+^. (**c**) *P. goldsteinii* lysate was fractionated using anion exchange chromatography. The presence of the antigen was tested *in vitro* similarly to (b). (**d**) *P. goldsteinii* lysate was fractionated using cation exchange chromatography. The presence of the antigen was tested *in vitro* similarly to (b). (**e**) Extracts derived from recombinant *E. coli* expressing the indicated candidate proteins were tested *in vitro* for the presence of the antigen, similarly to (b). Dot plots are representative of at least 3 independent experiments. Cells were gated on Via-probe^-^ CD4^+^ Vα2^+^ Vβ6^+^. (**f**) The graph shows the mean and S_D_ of one representative experiment described in (e) of at least 3 independent experiments. Each symbol represents one technical replicate. (**g-h**) Congenically marked WT CD45.1^+^ mice received 7X10^5^ CellTrace Violet-labeled naïve CD4^+^ TN T cells (CD45.2^+^), and were immunized sub-cutaneously with 25μg of bacterial extracts (g) or with 2 μg of peptide in alum (h) 1 day later. 3.5 days after immunization, proliferation was assessed in the draining lymph node (inguinal Lymph Node) by flow cytometry. The dot plots show dilution of CellTrace-Violet in CD45.1^-^CD45.2^+^CD4^+^Vα2^+^Vβ6^+^ T cells in one representative mouse per condition. Mutations in the core epitope are indicated in red. (**i**) The graph shows all mice analyzed in two independent experiments. Experiments involving immunization with bacterial extracts and peptides were done independently. The graph shows the means +/− SDs and each symbol represents a single mouse (Vector n=4, β-gal n=3, *P.d* β-hex n=5, *P.m* β-hex n=3, *P.g* β-hex n=7, *P.m/P.g* peptide n=5 each). (**j**) Same as (b), using as the source of antigen peptide concentrations of 500nM-50pM in serial 10-fold dilutions. The graph shows the mean and SD of one experiment, representative of at least three independent experiments. Each symbol represents one technical replicate. Cells were gated on Via-probe^-^CD4^+^Vα2^+^Vβ6^+^. Any of the activating peptides (*P.g*, *B.l* or *B.v*) compared to any of the inactive peptides (*P.n, B.o, P.d* or *P.m*) yield P values <0.0001 using 500nM peptide as a reference. (**k**) Alignment of sequences homologous to the TN epitope. Green positions indicate predicted TCR contact sites and blue the I-A^b^ anchor positions. “Activity” represents the ability of each peptide to activate the TN TCR *in vitro* (see j). (**l**) Congenically marked CD45.1 ^+^ WT recipients were treated with antibiotics (ABX) for 2 weeks. At day 16, mice received 2X10^5^ naïve CD45.2^+^ CD4^+^ TN T cells. The following day, the mice were colonized (*P. goldsteinii* or *B. vulgatus* n=8 each) or not (n=7) with the indicated bacteria. Four weeks post-colonization, the small intestine intraepithelial compartment (IEL) and the mesenteric lymph nodes (mLN) were harvested and analyzed by flow cytometry. (**m**) Dot plots show the frequency of TN cells in the IEL (top) and mLN (bottom) of one representative mouse per group. TN cells were gated on live Aqua^-^CD4^+^Vα2^+^Vβ6^+^CD45.2^+^CD45.1^-^CD8β^-/lo^. (**n**) Graphs show all mice analyzed in two independent experiments shown in (m). The frequency of CD4_IELs_ was assessed in the IEL (top) and the frequency of Tregs in the mLN (bottom). The graphs show the mean+/− standard deviation (SD) and each symbol represents a single mouse. P values were calculated using unpaired one-sided Student’s t tests. P values <0.05 were considered statistically significant. (*P.g= P. goldsteinii, B.l= B. luti, B.v= B. vulgatus, P.n= P. nanceiensis, B.o= B. ovatus, P.d= P. distasonis, P.m= P. merdae,* β-hex= β-N-acetylhexosamindase).

While the microbiome of the Whitehead Institute colony (WI) contained sufficient antigen to induce strong proliferation of the TN cells *in vivo*^6^, the relative abundance of the *P. goldsteinii* OTU was small compared to the microbiome of Taconic mice (Extended Data Fig. 3c). However, the WI microbiota was highly enriched in closely related species belonging to the *Bacteroidetes* phylum, one of the most abundant taxa in many animal facilities^35^. This prompted us to ask whether homologous sequences derived from related species could also activate the TN TCR which was originally cloned from a mouse housed at the WI facility. The corresponding β-hex peptides (non-N-acetylated forms) derived from *B. vulgatus* and *Bacteroides luti* could indeed induce proliferation of TN cells *in vitro,* but only at higher concentrations (Fig. 3j,k). Using a combination of BlastP and Jackhmmer analyses^36^, we found possible immunostimulatory sequences in a wide range of organisms belonging to the *Bacteroidetes* phylum, from *Bacteroides* and *Parabacteroides* genuses to more distantly related species such as *Spirosoma panaciterrae* (Extended Data Fig. 6). All such sequences mapped to the β-hex gene and over 99% of the species are found in the gastrointestinal tract. We also found several isolates containing a β-hex epitope among a library of 11,000 isolates derived from healthy human donors^37^, suggesting that the β-hex epitope recognized by TN T cells is also present in the human gut microbiota (Supplementary Data Table 3).

We then asked whether any of these bacteria could promote expansion and conversion of TN T cells into CD4_IELs_ and pTregs *in vivo.* WT SPF hosts were pre-treated with antibiotics and then received congenically-marked TN CD4^+^ T cells. Hosts were colonized with mouse isolates of *P. goldsteinii* or *B. vulgatus.* Colonization with either species promoted the expansion and development of CD4_IELs_ in the intestine and pTregs in the mLNs (Fig. 3l-n). We obtained similar results with immunodeficient mice (TCRαßKO) as recipients (Extended Data Fig. 7). The pTreg TN TCR therefore likely recognizes an even broader collection of bacterial species (encoding the ß-hex epitope) belonging to the *Bacteroidetes* phylum, rather than a single bacterial species. Microbes other than *P. goldsteinii* or *B. vulgatus* that express similar or even the identical antigen may thus be capable of engaging CD4^+^ T cells that would migrate to the intestine and convert into CD4_IELs_.

We then asked whether oral provision of antigen in microbiota-depleted mice would suffice to induce CD4_IELs_ as suggested in another model^5^. Prior to transfer of TN cells, we treated SPF recipient mice with broad-spectrum antibiotics to deplete the microbiota^31^. One day after T cell transfer, β-hex peptide was administered orally via gavage. Oral administration of peptide induced robust proliferation of TN cells in all gut-draining mLNs. In contrast, mice that received neither antibiotics nor peptide, therefore relying solely on indigenous commensals as a source of antigen, showed stronger proliferation in the jejunum/ileum mLN (Extended Data Fig. 8a-c). We conclude that the β-hex antigen is naturally presented in the distal parts of the small intestine. When the antigen is provided as a dietary component, it reaches all gut-draining LNs. This data is in agreement with compartmentalization of intestinal T cell responses by the gut segment drained by specific mLNs, as shown using dietary and pathobiont-specific models^38^. We also analyzed the fate of transferred TN cells four weeks later and found that the *P. goldsteinii* β-hex peptide induced CD4_IELs_ in the small intestine in the absence of microbiota (Extended Data Fig. 8d-g). Provision of this TCR ligand replaces the strict requirement of microbiota for CD4_IEL_ development in antibiotic-treated mice.

Next we sought to investigate whether this prevalent epitope that is able to induce CD4_IELs_ from TN precursors is also recognized by CD4_IELs_ of SPF mice. We designed a *P. goldsteinii* β-hex MHCII tetramer spanning the β-hex epitope, (Extended Data Fig. 9a) and enumerated β-hex specific CD4^+^ T cells by flow cytometry (Fig. 4a,b and Extended Data 9b). SPF mice showed few β-hex-positive cells in the mLNs (Extended Data Fig. 9b) compared to the IELs (Fig. 4a,b). Considering the diversity of TCRs present in mLNs^10^, this is not surprising. In the IE compartment, we observed an enrichment of *P. goldsteinii* β-hex specific CD4^+^ cells but not of the related *P. merdae* or *P. distasonis.* CD4^+^ T cells isolated from the IE compartment of GF mice did not bind to *P. goldsteinii* β-hex tetramer (Extended Data Fig. 9c). The frequency of β-hex-specific cells, both in mLNs and IEL varied in WT mice housed in different rooms (Fig. 4b and Extended Data Fig. 9b), potentially due to differences in microbiome composition. Accordingly, CD4_IELs_ specifically produce IFN-γ in response to *P. goldsteinii* β-hex peptide (Extended Data Fig. 9d,e). The *P. goldsteinii* β-hex epitope is therefore an abundant natural TCR ligand of intraepithelial CD4^+^ T cells.

**Figure 4:**
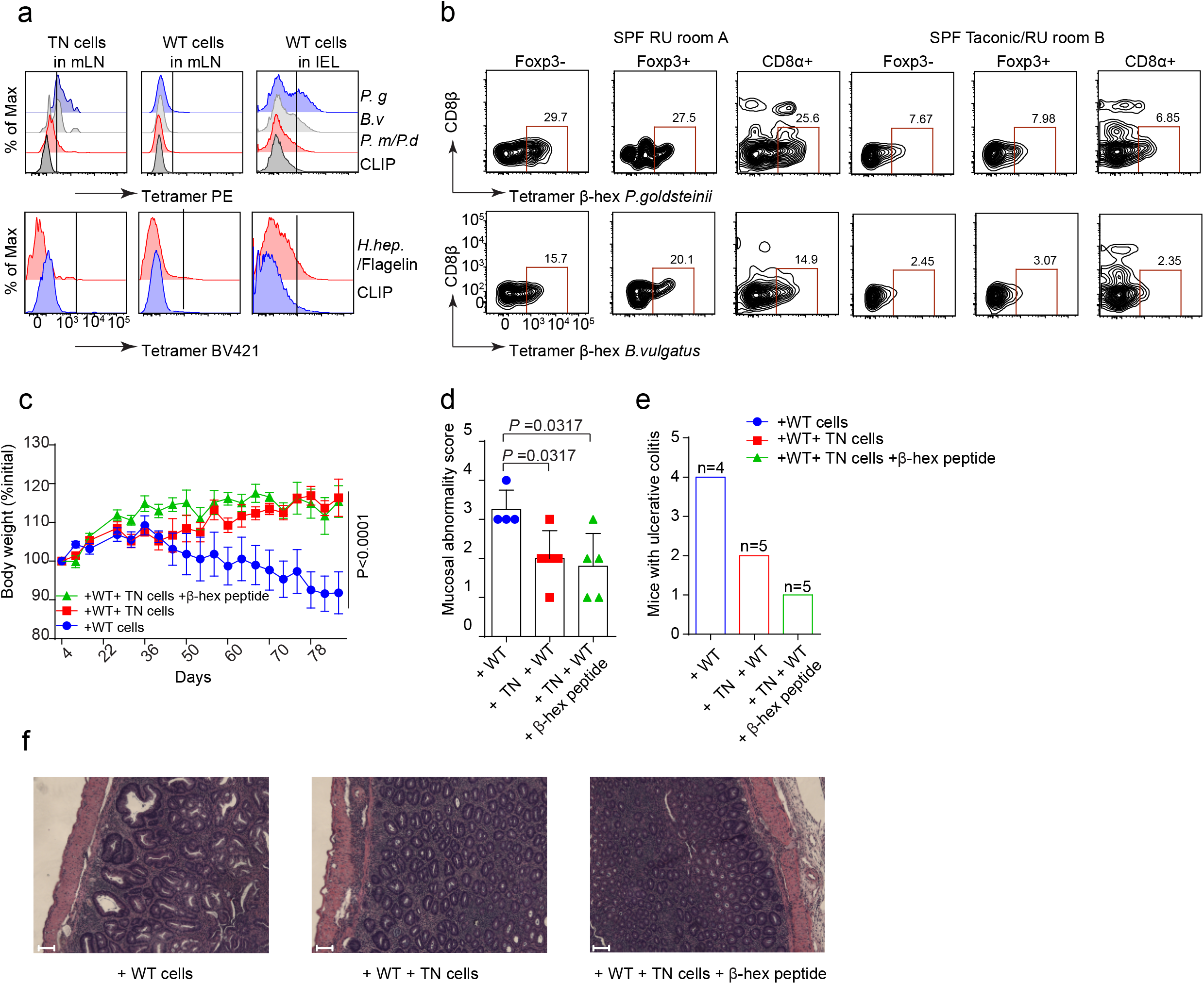
The β-hexosaminidase TN epitope is recognized by WT CD4IELs and mediates protection against intestinal inflammation. (**a**) Naïve TN cells were transferred 24h prior to tetramer staining into congenically-marked WT SPF hosts. Mesenteric lymph nodes (mLNs) and IELs were harvested and stained using the indicated tetramers. Cells were gated on live Aqua^-^ CD45^+^CD4^+^TCRβ^+^TCRγδ^-^CD8β^-/lo^. (a,b) Each Samples were processed separately for three mice and then pooled for tetramer staining. (**b**) Cells from the IEL of SPF mice housed at Rockefeller University (RU) either bred at RU for many generations (SPF RU room A) or bought from Taconic 6 months prior to analysis and housed at RU in a different room (SPF Taconic/RU room B) were stained with the indicated tetramers (experiment n=4). (**c**) Taconic Rag2KO received 2X10^5^ naïve CD45.2^+^CD4^+^ TN cells. Twelve days later, mice received 3X10^5^ naïve CD45.1^+^CD4^+^ WT cells (day 0). One group of mice received 3 doses of 500μg of *P. goldsteinii* β-hexosaminidase (β-hex) peptide (KWDYKGSRVWLNDQ) orally by gavage (at day −10, −8,1). Mice were weighed every other day throughout the experiment. The mice were euthanized either when they lost 20% of their initial weight or at the end of the experiment (day 81). The graph shows weight loss of the mice in the different groups. The graphs show mean/group +/− standard error. P values were calculated using a two-way ANOVA. P values <0.05 were considered statistically significant. (**d**) Mucosal abnormality score of colon H&E stained sections. Mucosal abnormality score is based on histology shown in (f). A Mann-Whitney test was used here (non-parametric data set). P values <0.05 were considered statistically significant. (**e**) Fraction of mice with colonic ulcers at the experimental end point, based on histology analysis shown in (f). (**f**) Photomicrographs of H&E stained large intestine (colon) sections of mice from the indicated colitis groups, representative of 5 mice/group. Scale bar=50μm.

Next we asked whether TN CD4_IELs_, similarly to Tregs, could protect against intestinal inflammation^39^. Forced conversion of CD4 cells into CD4_IELs_ protects against colitis^18^. Transfer of TN cells into immunodeficient recipients do not cause overt intestinal pathology, despite robust proliferation of the transferred cells and IFNγ production^6^. RNAseq analysis showed that TN CD4_IELs_ express a number of genes associated with regulatory functions^6^. To test the anti-inflammatory potential of TN CD4_IELs_, we first transferred naïve CD4^+^ TN T cells into RAG-deficient mice, and after 12 days we transferred congenically-marked WT naïve CD4^+^ T cells (Extended Data Fig. 10a). WT cells expanded in all organs analyzed, irrespective of the presence of TN cells (Extended Data Fig. 10b). TN cells converted into CD4_IELs_ 4-8 weeks post-transfer but did not differentiate into pTregs (Extended Data Fig. 10c,d), similarly to other commensal-specific T cells that fail to differentiate into pTregs when transferred into Rag-deficient hosts^40^. Mice that received both TN and WT cells with or without oral β-hex peptide administration were partially protected from colitis, as judged by reduced weight loss and improved histopathological score (Fig. 4c-f). The majority of mice that received TN cells were free of ulcers in the colon (Fig. 4e). The frequency of WT-derived Tregs was low in all groups, suggesting little or no contribution of pTregs to such protection (Extended Data Fig. 10c,d). When mice received both TN and WT cells, we observed a slight increase in the frequency of WT-Tregs in the LP. Therefore, we cannot formally rule out that those few pTregs contributed to the protective effect observed in mice transferred with TN cells. We conclude that, in the presence of their cognate antigen, TN CD4^+^ T cells can protect against intestinal inflammation.

Our novel commensal-specific model provides a new tool to study CD4_IELs_ development and can be used to understand how different commensal bacteria shape plasticity, intestinal adaptation and function of CD4_IELs_. The microbiota that resides in different niches of the intestinal mucosal surface can drive functional specialization of CD4^+^ T cells. We show here, that *P. goldsteinii* and potentially many other *Bacteroidetes* which reside in the lumen and possibly the mucus layer (e.g. *Parabacteroides goldsteinii*)^41^ are able to induce antigen-specific CD4_IELs_ in a two-step manner. First, by the engagement of the TCR with its cognate antigen, and second by the microbiome-modified intestinal epithelial environment^42^. Recognition of an antigen such as β-hex, present in a variety of abundant commensals *(Bacteroidetes)* might provide a competitive advantage to CD4_IEL_ precursors to reside in the intestinal epithelium. Deciphering the rules that govern the mutualism between commensals, pathobionts and immune cells shall help us better understand intestinal homeostasis and inflammation, with prospects for controlling inflammation.

## Author contributions

A.B., D.B. and H.P. conceived the study, A.B and D.B designed and performed the experiments. A.B, D.B and H.P wrote the manuscript. D.B performed the biochemical experiments to identify and characterize the TN antigen, with help from A.B and P.B. D.B and A.B. performed the *in vivo* experiments, with help from M.L and P.B. M.L. performed the characterization of WT CD4IELs (Extended Data Fig. 1). M.M supported D.B and A.B with anaerobic cultures, provided mouse isolates and experimental guidance. S.O. analyzed the 16s sequencing data. M.P. provided human bacterial isolates, sequenced and assembled the genomes of the mouse bacterial isolates used in this study. R.C provided support for peptide synthesis and J.L guided D.B with antigen fractionation. J.S. performed MHCII peptide binding assays, under the supervision of A.S. E.A and D.M commented on project design and assisted in data interpretation. H.P and A.B provided guidance and supervised the study. D.B performed the phylogenetic and statistical analyses.

## Acknowledgements

We thank S. Kolifrath and J.Jackson for technical support with mouse husbandry, genotyping and colony management at Harvard, and A. Rogoz and S. Gonzalez at Rockefeller University. We thank the hematology flow cytometry facility of BCH, in particular R. Mathieu for assistance with cell sorting, and K. Gordon and K. Chhophel at Rockefeller. E. Spooner from the WIBR proteomics core for mass spectrometry analysis, R. Bronson from the rodent histopathology core of Harvard Medical school for histology and scoring. We thank the NIH tetramer core facility for providing all tetramers used in this study, in particular R.Willis for help with staining troubleshooting. We thank T.Lu and P.Silver for allowing us to use their anaerobic chambers and J.Leube for help with the initial *in vitro* experiments. We are in indebt to C.McClune for assistance with cloning, bioinformatic analyses and project suggestions and D. VanInsberghe for support with phylogenetic analysis. We are grateful to G.Victora for invaluable guidance and support throughout this project. We thank A.Woodham for assistance with ELIspot analysis and all the members of the Ploegh lab for suggestions and fruitful discussions. We thank G. Victora, B. Reis and G. Donaldson for critical discussions and suggestions. This study was initiated at the Whitehead Institute for Biomedical Research.

## Supplementary Information

The Supplemental Data file contains the Supplementary Tables 1-3 as well as 16s analyses, primers and isolates used in this study.

## Data availability

Raw data for all figures is provided with the paper. Sequencing data is provided in Supplementary Data file.

## Materials and methods

### Mice

Mice were housed at the Whitehead Institute for Biomedical Research (WIBR), Boston Children’s hospital (BCH) and Rockefeller University under specific pathogen–free conditions, in accordance with institutional guidelines and approved by the institutional animal care and use committee of the Massachusetts Institute of Technology, the Boston Children’s Hospital (IACUC protocol number 16-12-3328) and Rockefeller University. pTreg TN mice were generated as described^1^. C57BL/6, C57BL/6 CD45.1, Rag1-/-, Rag2-/-, TCRα-/-, TCRβ-/-, mice were purchased from Taconic Biosciences and Jackson Laboratory and maintained at our facilities. pTreg TN were crossed to C57BL/6 RAG1KO for at least eight generations to generate pTreg TNxRAGKO (TN). TCRαβKO (TCRα^-/-^β^-/-^) were generated by intercrossing TCRα^-/-^ and TCRβ ^-/-^. Germ-free (GF) C57BL/6 and Swiss Webster mice were kept in germ-free flexible film isolators (Class Biologically Clean Ltd) at Rockefeller University. For colonization experiments, GF C57BL/6 mice were exported to isocages bioexclusion system (Tecniplast) and housed in isocages for the duration of the experiment (3-5 weeks as indicated in figure legends). Mice housed in tecniplast were given autoclaved food and water.

### Antibodies, reagents, and FACS analysis

Fluorescently labeled antibodies were purchased from BD bioscience (anti-Vβ6, RR4-7; anti-CD3, 145-2C11; anti-Vβ5, MR9-4), eBioscience (anti-CD45.1, A20; anti-Foxp3, FJK-16s; anti-CD4, RM4-5; Va2, B20.1; anti-CD11c, N418). BioLegend (anti-CD45.2, 104; anti-CD8α, 53-6.7; anti-CD8β, YTS 156.7.7; anti-CD44, IM7; anti-I-A/I-E, M5/114.15.2, α-CD3, 145-2C11). Proliferation was measured by dilution of cell proliferation dyes CFSE (Sigma-Aldrich) and CellTrace Violet (Life Technologies), according to manufacturer’s instructions. The cell viability dye 7-AAD (Via-Probe) and Live/dead fixable Aqua fluorescent reactive dye were purchased from BD Biosciences and ThermoFisher, respectively. Staining with live/dead markers was performed following manufacturer’s instructions. Staining for cell surface markers was performed at 4°C for 30min in PBS+1mM EDTA+ 0.5% bovine serum albumin. Intracellular staining was performed according to manufacturer’s instructions, using the Foxp3/Transcription Factor Staining Buffer Set (eBioscience). Flow cytometry data were acquired on an LSRFortessa (Becton Dickinson) instrument and analyzed with the FlowJo software package (Tri-Star). Alum (Imject) used for immunizations was obtained from Thermo Fisher Scientific.

### Antibiotic treatment

An antibiotic cocktail containing Vancomycin (0.5 g/liter), ampicillin (1.0 g/liter), neomycin (1.0 g/liter) (Amresco), and metronidazole (1.0 g/liter) (Sigma) was given ad libitum in the drinking water^2^ for the indicated periods as described in the figure legends.

### Bacterial growth and culture conditions

*Parabacteroides spp*. and *Bacteroides spp.* were grown on ASF plates (Becton Dickinson). Single colonies were picked and grown in an anaerobic chamber (Coy Lab Products) using 2.5% H_2_, 5% CO_2_ and 92.5%N_2_ in Brain Heart Infusion (BHI, Becton Dickinson) broth supplemented with 5g/L yeast extract (Becton Dickinson), 10mg/L hemin, 1mg/L Vitamin K3/menadione and 0.5g/L of cysteine-HCl. Under anaerobic growth, both plates and liquid cultures were pre-reduced for 6-10h prior to inoculation. *E.faecalis* and *L.plantarum* were grown anaerobically in BHI and MRS broth, respectively. E.coli was grown aerobically in Luria Broth (LB).

### Isolation of bacterial strains

For isolates obtained from mice, 2-3 fecal pellets were homogenized in 1mL of sterile PBS with a stainless-steel bead in a Qiagen Tissue Lyser II. Samples were cleared for 30 sec at 500xg to remove large fecal debris. Supernatants were plated on the indicated media (see supplemental data) and incubated anaerobically for 48h at 37°C. Isolates were re-streaked twice for isolation of single colonies on the appropriate media before glycerol stocks were made. The supplemental data provide a detailed list of the different strains used in this study. For colonization experiments, *P.goldsteinii* isolates 841 and 33-2-9-2-8 were used in combination. Isolate 33-9-2-8 was used exclusively for antigen purification and as a source of DNA to build the candidate *E.coli* recombinants.

### Antigen purification

Overnight cultures of *P. goldsteinii* isolate 33-9-2-8 were diluted 1:100 and grown for another 24h under anaerobic conditions. Cultures were harvested at 2500g for 15min at 4°C, resuspended in 50mM Tris-HCl, pH 7.4 containing a protease inhibitor cocktail (1 tablet/50ml, Complete, Roche) and 1mg/ml of deoxyribonuclease I (Grade II, Roche) and lysed using a French press (FA-032 Standard CELL, Thermo Fisher). Proteins were precipitated from the lysate, cleared by centrifugation, using increasing concentrations of ammonium sulfate (Molecular biology grade, Sigma Aldrich) at 4°C for 15min and recovered at 10,000g for 15min. Precipitates were resuspended in 50mM Tris-HCl (pH 7.4) and dialyzed overnight (ThermoFischer, Slide-A-Lyzer™ Dialysis Cassettes, 3.5K MWCO). Activity of each fraction was measured in the TN T cell proliferation assay. Samples containing activity were fractionated by anion exchange chromatography on a Mono Q 5/200 GL column (GE Healthcare, Piscataway NJ) using a gradient from 50mM Tris-HCl (pH 7.4) to 50mM Tris-HCl (pH 7.4) 500mM NaCl. To collect sufficient material, this step was repeated 5 times. Fractions that yielded proliferation of TN T cells were pooled and subjected to cation exchange chromatography on a Mono S 5/200 GL column (GE Healthcare, Piscataway NJ). The active fractions were again pooled and subjected to buffer exchange (Amicon ultra 3k cutoff, sigma Millipore).

### Mass spectrometry analysis

Active fractions from the MonoS chromoatography step were recovered by TCA precipitation, resuspended in a Tris/ Urea buffer, reduced with DTT, alkylated with iodoacetamide and digested with 2μg trypsin/100μl of sample at 37°C overnight. The resulting peptides were washed, extracted and concentrated by solid phase extraction using Waters Sep-Pak Plus C18 cartridges. Organic solvent was removed and the volumes were reduced to 15 μl using a speed vac for subsequent analyses. The sample was then injected onto a Waters NanoAcquity HPLC equipped with a Aeris 3 μm C18 analytical column (0.075 mm by 20 cm; Phenomenex). Peptides were eluted using standard reverse-phase gradients. The effluent from the column was analyzed using a Thermo Orbitrap Elite mass spectrometer (nanospray configuration) operated in a data-dependent manner for the duration of the 90 minute run. The resulting fragmentation spectra were matched against the Refseq entries for *P.goldsteinii* using Mascot (Matrix Science). Scaffold Q+S (Proteome Software) was used to provide consensus reports for the identified proteins.

### Isolation of mononuclear cells from intestine

Small intestines were harvested and mononuclear cells were isolated from the IE compartment and lamina propria as described^1,3^.

### Tetramer staining

Cells isolated from the organs indicated in figure 4 and figure S9 were incubated with an Fc blocking agent (BD Pharminogen) for 10min at 4°C. Cells were then stained with tetramers (14 μg/mL in PBS + 0.5% BSA + 1mM EDTA + 0.05% Sodium azide) at 37°C for 1 h45min. Cells were counterstained for surface and intracellular markers as described above. For each experiment, 2-3 mice were processed separately and the cells obtained from them pooled for staining with the indicated tetramers.

### Elispot

IFNγ ELISpot assays were performed following the manufacturer’s recommendations (BD ELISPOT Mouse IFNγ ELISPOT Set, BD Biosciences). Briefly, 96-well ELISpot plates were coated with an IFNγ capture antibody in PBS overnight at 4°C. The plates were blocked in complete RPMI for 2h at RT. CD4_IELs_ (CD45^+^CD4^+^Aqua^-^TCRβ^+^TCRγδ^-^CD8β^-^ CD8α^+^) and CD4^+^CD8α^-^ (CD45^+^CD4^+^Aqua^-^TCRβ^+^TCRγδ^-^CD8β^-^CD8α^-^) isolated from the IEL of 3-5 Taconic SPF mice (4-9 months old) were sorted and co-cultured with DCs purified from the spleens of TCRαβKO mice, as described below (*in vitro* proliferation assay). 20,000-35,000 T cells were combined with DCs in a 1:2 ratio in duplicate in the presence or absence of 1μM of the indicated peptides. Positive control wells contained 1x Cell Stimulation Cocktail (Invitrogen) or 100nM α-CD3 (Biolegend). After plating, cells were incubated for 18h at 37°C. Plates were then washed and incubated with detection biotinylated anti-IFNγ antibody for 2h and followed by incubation with streptavidin-horse radish peroxidase (BD Biosciences) for 1h. Plates were developed with 3-amino-9-ethyl-carbazole substrate (BD ELISPOT AEC Substrate Set) for 10 minutes and air-dried for at least 18 hours. Spots were enumerated using an ImmunoSpot Ultimate Analyzer (Cellular Technology Limited) and normalized to the number of cells seeded to enable comparisons between experiments.

### qPCR of fecal DNA

DNA was extracted from fecal pellets using the Qiagen power fecal kit, following the manufacturer’s instructions. qPCR was performed using SYBR green master mix (BioRad). Primers used for 16s were 5’-gtgStgcaYggYtgtcgtca-3, and 5’-acgtcRtccMcaccttcctc-3^4^ 5’-gcagcacgatgtagcaataca-3’ and 5’-ttaacaaatatttccatgtggaac-3’ for *P. goldsteinii*^5^ and 5’-atagcctttcgaaagraagat-3’ and 5’-ccagtatcaactgcaatttta-3’ for *Bacteroides*^6^. The relative abundance of *P. goldsteinii* and *Bacteroides* were calculated by normalizing bacteria-specific CT to total 16s CT (i.e Relative Abundance of *P. goldsteinii* = 2^-(CT *P. goldsteinii* -CT 16s^)).

### Fecal bacterial growth

Fecal bacteria from the indicated mice were harvested by resuspending single fecal pellets into 500μl of PBS and spinning briefly to remove fibers. Supernatants were diluted 1:10, 1:100 and 1:1000 and then streaked on Schaedler blood agar, Columbia CNA agar or Bacteroides Bile Esculin Agar (BBE) (Becton Dickinson). Bacteria were grown for 48-72h aerobically or anaerobically as indicated. For the isolation of aerotolerant extracts, ~10 colonies from an anaerobic Schaedler blood agar plate were re-streaked to obtain single colonies, grown for 72h and then exposed to aerobic conditions (stored in the fridge outside anaerobic chamber). The plates were then placed back in the anarerobic chamber and incubated for further 72h before isolation of regrown bacteria.

### Bacterial extracts for proliferation assays

Unless specified otherwise, bacterial extracts were prepared from overnight cultures or from bacterial colonies scraped directly off agar plates into PBS. Bacterial pellets obtained by centrifugation at 13,500g for 10 min at 4°C were resuspended in 1 ml of PBS containing protease inhibitors (cOmplete Mini, Roche; 1 tablet per 10 ml of PBS following the manufacturer’s instructions) and deoxyribonuclease I (grade II, Roche). Bacteria were lysed by three rounds of sonication (Branson Sonifier 450) for 1 min, 60 pulses, followed by 5 min of cooling on ice between each round (output control, 4; duty cycle, 50%). Cellular debris was removed by centrifugation at 14,000g for 30 min at 4°C. Protein concentrations were assessed using a bicinchoninic acid assay (BCA) assay. For *P. golsteinii* and *P. distasonis* extracts used for the *in vivo* proliferation assays, overnight cultures were spun down at 2,500g for 15min at 4°C, resuspended in PBS containing protease inhibitor cocktail (1 tablet/50ml, Complete, Roche) and 1mg/ml of deoxyribonuclease I (Grade II, Roche) and lysed using a French press (FA-032 Standard CELL, Thermo Fisher). Supernatants were cleared for 30min at 4,000g in a 100k Amicon column (Sigma-Millipore), followed by two PBS washes. The retentate was used for *in vivo* proliferation assays. Protein concentrations were determined using a BCA assay.

### Cloning recombinant *E. coli* and extract preparation

The genes of candidate proteins identified in the LC-MS/MS analysis were cloned into Pet-30b+ (EMD biosciences) using a Gibson cloning strategy. The inserts were designed to contain a C-terminal HA tag to allow confirmation of expression by immunoblot. After introduction into competent BL21 cells candidate antigens were expressed overnight in Terrific Broth at 30°C upon isopropyl ß-D-thiogalactopyranoside induction (1 mM) at an OD600 of ~0.6. Overnight cultures were harvested at 13,500g for 10 min at 4°C. The bacterial pellet was resuspended in 1 ml of PBS containing protease inhibitors (cOmplete Mini, Roche; 1 tablet per 10 ml of PBS following the manufacturer’s instructions) and deoxyribonuclease I (grade II, Roche). Extracts were then prepared for *in vitro* and *in vivo* proliferation as described above. For each candidate protein, several independent clones were tested in an *in vitro* proliferation assay (see above).

### MHC purification and binding assays

Purification of H-2 I-A^b^ and I-A^d^ class II MHC molecules by affinity chromatography, and the performance of peptide binding assays based on competition for binding of a high affinity radiolabeled peptide were performed as detailed elsewhere^7^. Briefly, the mouse B cell lymphoma LB27.4 was used as a source of Class II MHC molecules. A high affinity radiolabeled peptide (0.1-1 nM; peptide ROIV, sequence YAHAAHAAHAAHAAHAA) was co-incubated at room temperature with purified Class II MHC in the presence of a cocktail of protease inhibitors and a candidate inhibitor peptide. Following a two-day incubation, Class II MHC-bound radioactivity was determined by capturing MHC/peptide complexes on mAb (Y3JP, I-Ab; MKD6, I-Ad) coated Lumitrac 600 plates (Greiner Bio-one, Frickenhausen, Germany), and measuring bound radioactivity using the TopCount (Packard Instrument Co., Meriden, CT) microscintillation counter. The concentration of peptide yielding 50% inhibition of the binding of the radiolabeled peptide was calculated. Under the conditions used, where [label]<[MHC] and IC50 ≥ [MHC], the measured IC50 values are reasonable approximations of the true K_D_ values^8,9^. Each competitor peptide was tested at six different concentrations covering a 100,000-fold range, and in three or more independent experiments. As a positive control, the unlabeled version of the radiolabeled probe was included in each experiment.

### *In vitro* proliferation assay

Dendritic cells were purified from spleens and mesenteric lymph nodes of B16-FLT3L-injected mice using the Pan Dendritic Cell Isolation Kit (Miltenyi, Germany). Naïve CD4^+^ T cells were purified from the spleen and mesenteric lymph nodes of pTreg TN/Rag1 KO mice following the manufacturer’s instructions (Miltenyi, Germany) and labeled with 2.5 μM CFSE (Sigma) or 3μM CellTrace Violet (ThermoFischer). Purity was always greater than 98% for both DCs and T cells. 1X10^5^ DCs were incubated with bacterial extract (50μg/ml) or peptide (500nM, 50nM, 5nM, 0.5nM, 500pM, 50pM) for 2h before adding 1X10^5^ T cells. Proliferation was assessed by dilution of CFSE or CellTrace Violet after 3.5 days of co-culture.

### *In vivo* proliferation assay

TCRαβKO or CD45.1 mice received the indicated number of CellTrace Violet or CFSE-labeled CD4^+^ T cells isolated from pTreg TN/Rag1KO mice. Recipients were then immunized with 25μg of bacterial lysate in alum sub-cutaneously or injected intranasally with 2μg of peptide in PBS (no alum) 24h later. Proliferation of cells isolated from the draining lymph node (inguinal lymph node or mediastinal lymph node) was assessed by dilution of the cell proliferation dye.

### Colonization

For colonization of GF mice with *P.goldsteinii,* bacteria from frozen glycerol stocks were grown anaerobically for 48h on Schaedler blood agar plates. Single colonies were inoculated in BHIS supplemented with hemin, vitamin K3 and cysteine (see above) and cultures were harvested 18h later. Pelleted bacteria were resuspended in PBS (5ml of culture resuspended in 500μl of PBS) and given orally to GF mice by gavage (200μl/mouse). WT Jax mice were pre-treated with broad spectrum antibiotics for 1-2 weeks prior to colonization. Colonization was performed as described for GF mice above but mice received bacteria orally for three consecutive days. Colonization status was confirmed by qPCR or 16S sequencing.

### Peptide synthesis

Peptides were synthesized on a rink-amide linker resin to produce C-terminal amides (Advanced ChemTech) using a flow-based solid-phase peptide synthesizer at 60°C, as described^10^. Deprotection steps were performed by incubating the resin for 45 sec in dimethylformamide (DMF) with piperidine (20% vol/vol) using a constant flow of 20ml/min. The resin was then rinsed at the same flow rate for 1.5 min with DMF. Coupling was performed using standard Fmoc-protected amino acids (4 equivalents), N,N,N,N-Tetramethyl-O-(1H-benzotriazol-1-yl)uronium hexafluorophosphate (HBTU, 4 equivalents), and diisopropylethylamine (DIPEA, 8 equivalents) in DMF at a flow rate of approximately 3 mL per minute. At the end of the synthesis, the resin was rinsed in methanol, followed by dichloromethane and air-dried. Peptides used for *in vivo* experiments were acetylated at the N-terminal amine to increase their stability by inhibiting proteolysis at the N-terminus. N-terminal acetylation was performed on resin by incubation of N-terminally deprotected peptides with a mixture of acetic anhydride, DIPEA and DMF (1:2:8 vol/vol) for 10 minutes, followed by washing with DMF. Peptides were cleaved off the resin and deprotected using 92.5% trifluoroacetic acid, 5% H2O, and 2.5% TIPS, precipitated using cold diethyl ether, collected by centrifugation, air-dried, purified using reversed-phase C18 HPLC (Shimadzu), and lyophilized. Identity of the peptides was confirmed by liquid chromatography-mass spectrometry (Waters Xevo system). Purified peptides were resuspended in deionized water at a concentration of 1mg/ml and stored at −20°C. Peptides for oral gavage were ordered on Genscript at a purity >= 85%. Minimal epitopes used in *in vitro* proliferation assays were ordered from Genscript (≥ 95% pure).

### Isolation of CD4^+^ T cells

Naïve CD4^+^ T cells were isolated from spleen and mesenteric lymph nodes using the naïve CD4 T cells isolation kit (Miltenyi, Germany). Purity was always >98%.

### Histopathological analysis

Small and large intestines were fixed in 10% (v/v) buffered formalin, paraffin-embedded and stained with hematoxylin and eosin (H&E). Slides were scored blindly by a pathologist at the rodent histopathology core of Harvard Medical School.

### Statistical analysis

Mean and standard deviation/error were calculated using GraphPad Prism (GraphPad Software). Unpaired one-sided Student’s t tests were used to compare two variables. A Mann-Whitney test was used for non-parametric data sets. Two-way ANOVA was used for analysis change in weight loss in the colitis experiment. P values <0.05 were considered significant and where applicable are indicated in each figure legend.

### 16S analysis

DNA was extracted from fecal pellets using Quick-DNA Fecal/Soil Microbe Miniprep Kit (Zymo Research), following the manufacturer’s instructions. Paired-end 16s Illumina sequencing libraries were constructed using nested PCRs as described^11^. Libraries were multiplexed and sequenced on an Illumina MiSeq using paired-end 150bp reads at the BioMicro Center of MIT. Primers were trimmed, allowing at most 2 mismatches in the primer sequence. Forward and reverse reads were merged, allowing at most 2 mismatches in the merged sequence and sequences of length 253 ± 5 nucleotides, and quality filtered, discarding sequences with at most 2 expected errors. 99% OTUs were identified with UPARSE^12^ OTUs were assigned to genera using the Ribosomal Database Project naïve Bayesian classifier^13^ with 80% bootstrap confidence.

---

**Extended Data Fig. 1:**
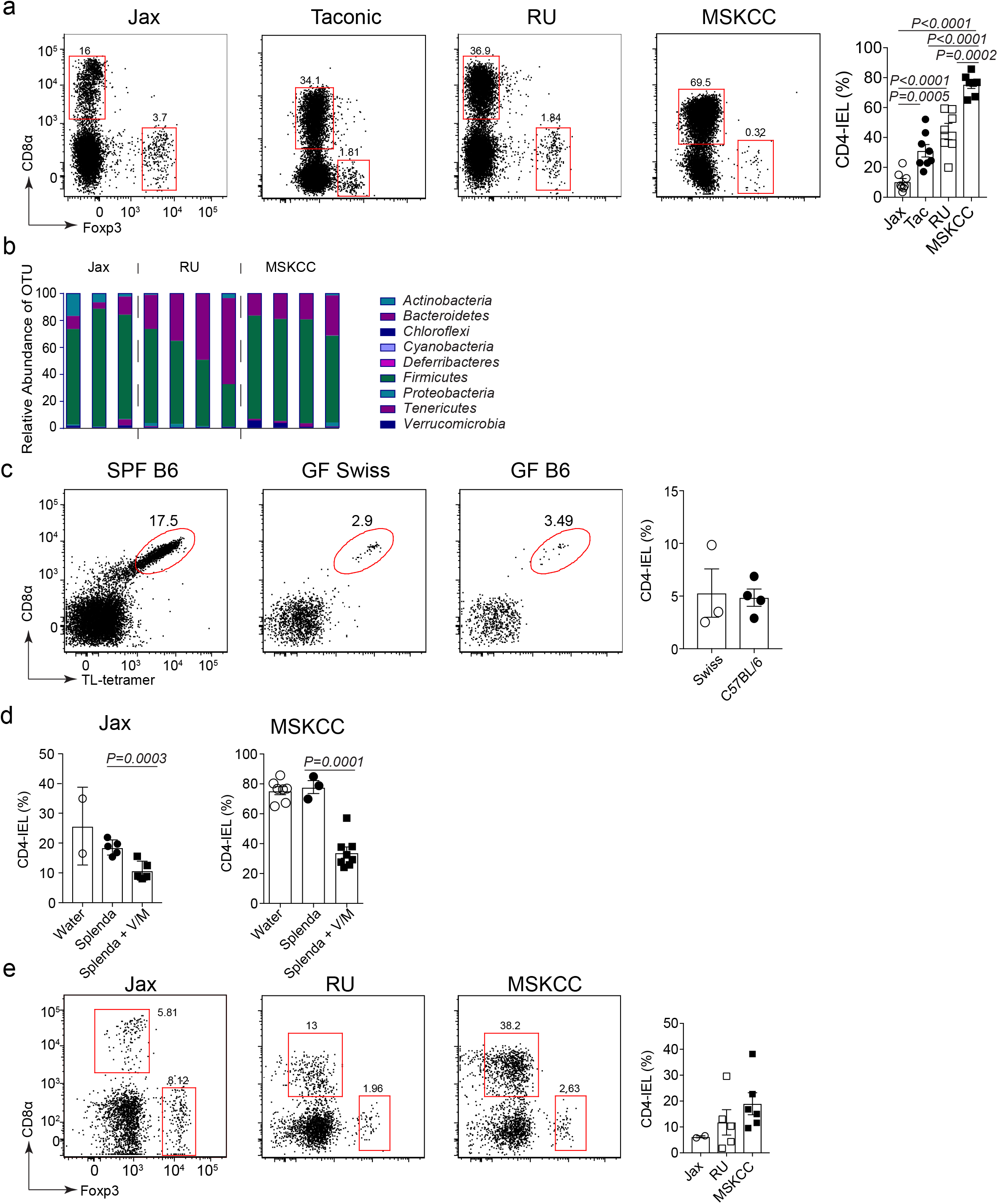
The microbiome influences the development and maintenance of CD4_IELs_. (**a**) Frequency of CD4_IELs_ in mice housed in different facilities. CD4_IELs_ from 8-16-week-old WT mice were harvested and their frequency determined by flow cytometry. (left) Dot plots show one representative mouse for the different experiments performed on different days (Jax n=8, Tac n=8, RU =7, MSKCC n=7). Graph on the right shows the quantitation of all mice analyzed. (**b**) The graph shows the relative abundance of the indicated phyla in WT mice housed in 3 different facilities (Jax=3, RU=4, MSKCC=4). Total DNA was extracted from fecal pellets upon arrival at the RU facility. The V4 region of 16s rRNA was amplified by nested PCRs and sequenced using illumina technology. The graph shows all mice analyzed in two experiments. (**c**) Frequency of CD4_IELs_ in germ-free (GF) mice, analyzed after export from their isolator. (Swiss Webster GF n=3, C57BL/6 GF n=4). (left) Dot plots of one representative mouse per group. (Right) Quantitation of all mice analyzed. (**d**) Frequency of CD4_IELs_ in eleven-week-old Jax mice treated with vancomycin and metronidazole (V/M) for 8 weeks (left) and in five-week-old mice from MSKCC treated with V/M for 4 weeks (right). (V/M) = 0.5g/L vancomycin /1g/L metronidazole. This combination was chosen because it impacted CD4_IELs_ without affecting the frequency of Tregs in the IE compartment. (Jax water n=2, Jax Splenda n=5, Jax Splenda + V/M n=5, MSKCC water n=7, MSKCC Splenda n=3, MSKCC Splenda + V/M n=). (**e**) Fecal matter transplant (FMT) from the indicated mice into GF mice. Four weeks after FMT, the frequency of CD4_IELs_ was assessed by flow cytometry. Dot plots show one representative mouse/group. The graph on the right shows all mice analyzed. (Jax n=2, RU n=5, MSKCC n=6). (a,c,e) Dot plots shows cells pre-gated on CD45^+^CD8β^-^CD4^+^TCRβ^+^TCRγδ^-^. (Jax= Jackson, Tac= Taconic, RU=Rockefeller University, MSKCC= Memorial Sloan Kettering Cancer Center). P values were calculated using unpaired one-sided Student’s t tests. P values <0.05 were considered statistically significant. The graphs show the mean and standard deviations. Each symbol represents a single mouse.

**Extended Data Fig. 2:**
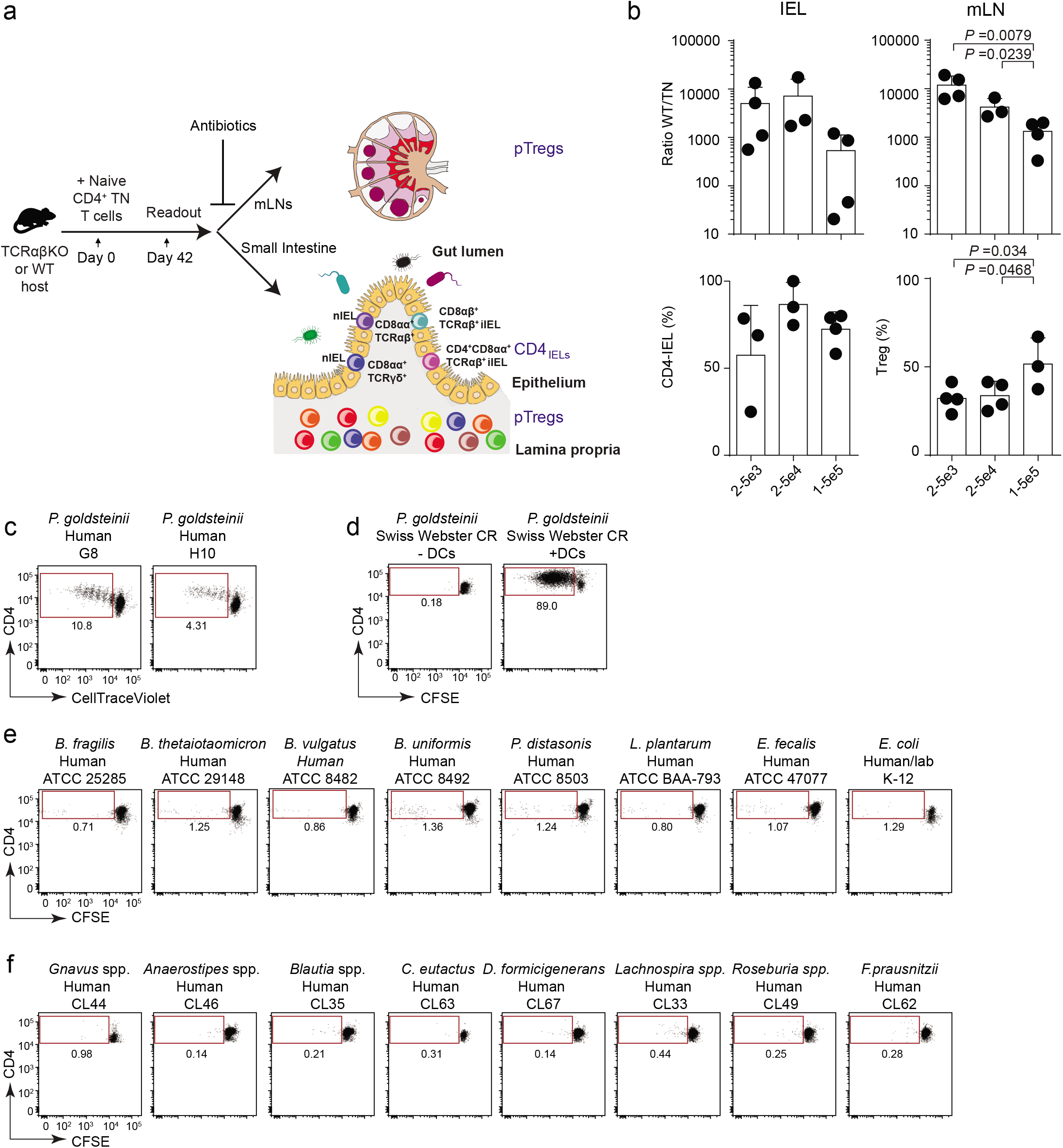
The pTreg TN TCR recognizes the predominant member of the Altered Schaedler Flora. (**a**) Schematic of the TN model. (**b**) Congenically-marked WT hosts received different numbers of naïve CD4^+^ TN T cells (2-5X10^3^ n=4, 2-5X10^4^ n=3, 1-5X10^5^ n=4). After 4 weeks, the IEL and mesenteric lymph nodes (mLN) were harvested, stained and analyzed by flow cytometry. The graphs on top show the ratio of WT/TN cells in the IEL (left) and mLN (right), respectively. The bottom graphs show the frequency of CD4_IELs_ among TN cells in the IEL (left) and Tregs in the mLN (right). All graphs show the mean and standard deviation (SD); each symbol represents a single mouse. TN cells were gated on CD45.2^+^CD45.1^-^Aqua^-^ CD4^+^Vα2^+^Vβ6^+^ TCRβ^+^TCRγδ^-^ in addition to CD8β^-^CD8α^+^Foxp3^-^ for CD4_IELs_ and CD8β^-^CD8α^-^Foxp3^+^ for Tregs. WT cells were gated on CD45.2^-^CD45.1^+^Aqua^-^. (**c**) Naïve CD4^+^ TN T cells were labeled with CellTrace-Violet and co-cultured with dendritic cells (DCs) purified from B16-Flt3L-injected mice in the presence of bacterial extracts derived from the indicated bacteria isolated from healthy human volunteers. 3.5 days later, CFSE dilution was assessed by flow cytometry. Dot plots are representative of at least 3 experiments. Cells were gated on Via-probe^-^ CD4^+^ Vα2^+^ Vβ6^+^. (**d**) Same experimental setup as (c) but using CFSE instead of CellTrace-Violet and extracts derived from mouse *P. goldsteinii* in the presence or absence of DCs. (**e**) Same as (c) using human isolates bought from ATCC. (**f**) Same as (c).

**Extended Data Fig. 3:**
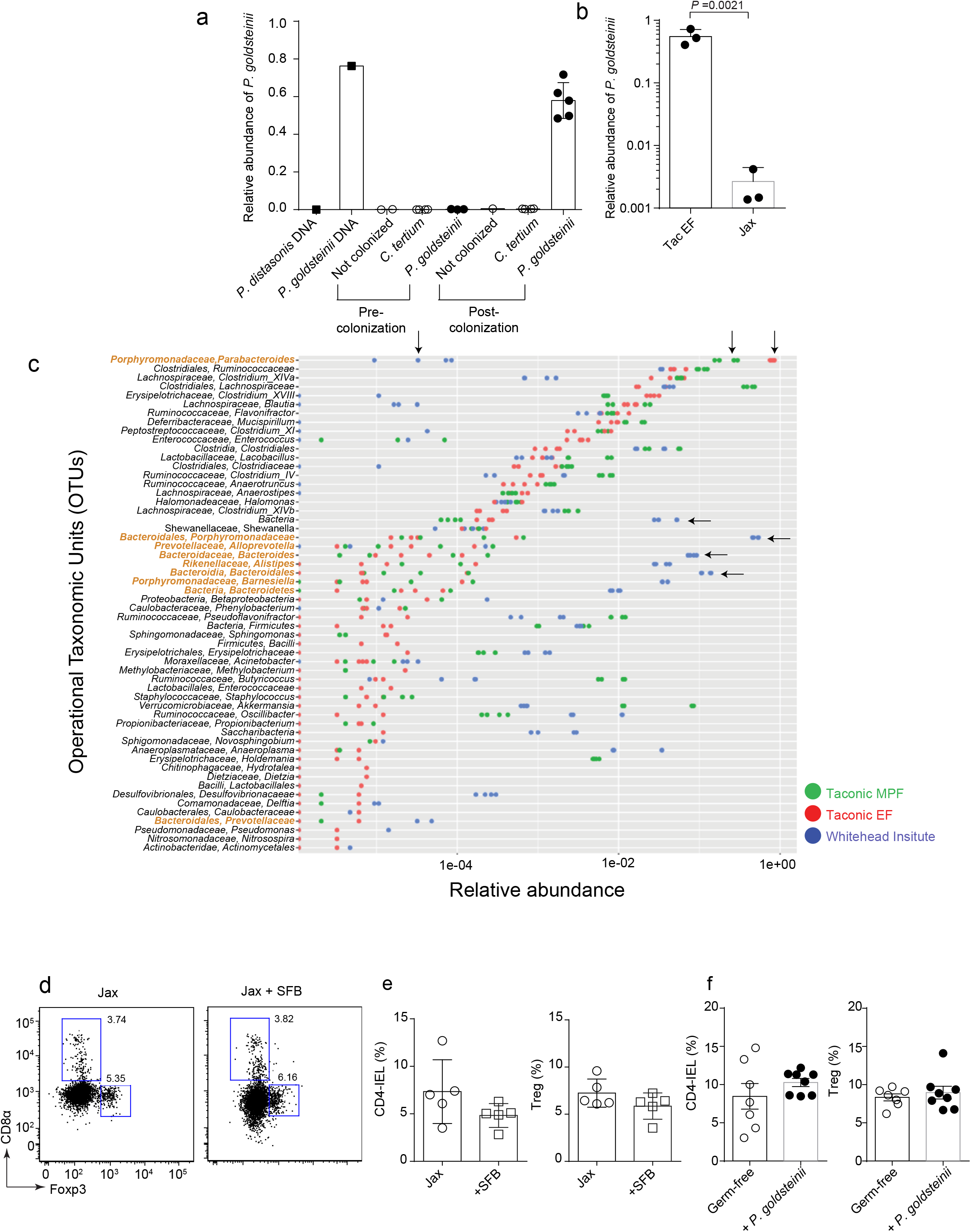
*P. goldsteinii* is abundant in SPF mice but does not rescue CD4IEL accumulation in germ-free mice. (**a**) At the experimental end point (~4 weeks postcolonization), total fecal DNA was extracted from Rag1 KO mice colonized or not with the indicated bacteria (see Fig. 2a-d). qPCR was performed using *P. goldsteinii-specific* and 16s generic primers. The graph shows the relative abundance of *P. goldsteinii* normalized to the total 16s (i.e Relative Abundance of *P. goldsteinii* = 2^-(CT *P. goldsteinii*-CT 16s)^) in all mice analyzed in two independent experiments. Each graph denotes the mean +/− SD and each symbol represents a single mouse. (**b**) Relative abundance of *P. goldsteinii* in the feces of Jackson (Jax) or EF mice as measured in (a) in all mice analyzed in one experiment (n=3/group). (**c**) The graph shows bacterial composition of fecal matter from mice housed in 3 different facilities. Total DNA was extracted from fecal pellets upon arrival at the WIBR facility (Taconic health status excluded flora (EF) and Taconic MPF) or in WT mice housed at WIBR facility. The V4 region of 16s rRNA was amplified by nested PCRs and sequenced using Illumina technology. Each symbol represents a single mouse in technical duplicate (n=2/facility). Arrows highlight the differences in bacterial composition between the flora analyzed. OTUs colored in orange are derived from *Bacteroidetes.* (**d**) Jax mice were colonized or not by gavage with SFB. After 4 weeks, IELs were harvested and stained for flow cytometry analysis. Dot plots show one representative mouse per group (n=5 for each group) of 1 experiment. (**e**) Graphs show all mice analyzed in (d). Graphs show the mean +/− standard deviation (SD) of all mice analyzed. (**f**) C57BL/6 germ-free (GF) mice (8-10 weeks-old), kept in isolator cages under GF conditions, were colonized or not (n=7) by gavage with *P. goldsteinii* (n=8) grown anaerobically in BHIS. After 3-5 weeks, IELs were harvested and stained for analysis by flow cytometry. Graphs show the mean +/− standard deviation (SD) of all mice analyzed in 3 independent experiments. P values were calculated using unpaired one-sided Student’s t tests. P values <0.05 were considered statistically significant.

**Extended Data Fig. 4:**
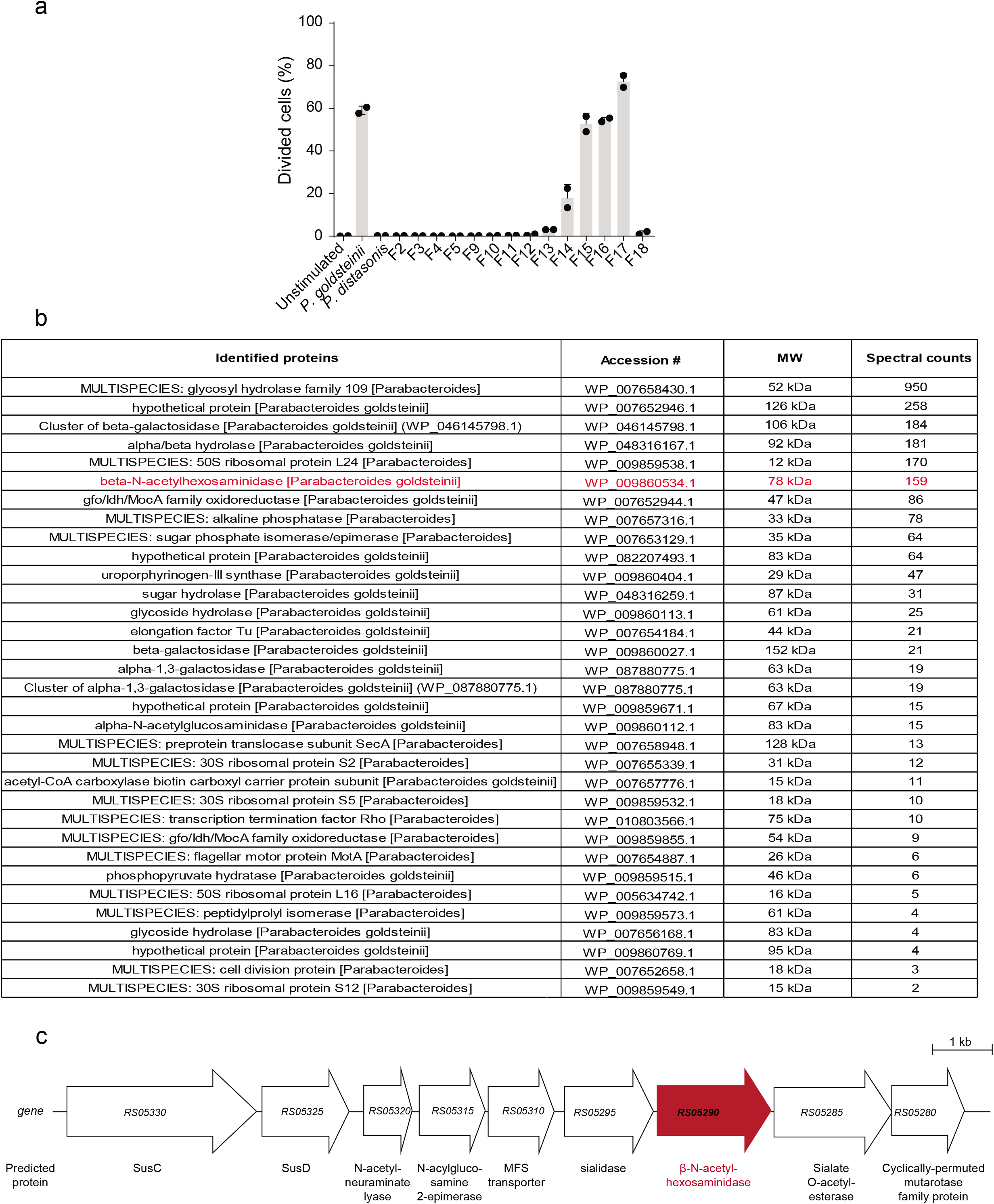
Mass spectrometry of *P. goldsteinii* antigen-enriched fraction. (**a**) *P. goldsteinii* lysate was fractionated using a combination of ammonium sulfate precipitation, anion and cation exchange chromatography. The resulting fractions were tested in an *in vitro* proliferation assay: Naïve CD4^+^ TN T cells were labeled with CellTrace-Violet and co-cultured with dendritic cells purified from B16-Flt3L-injected mice in the presence of the indicated fractions. 3.5 days later, CellTrace-Violet dilution was assessed by flow cytometry. The graph shows the mean and standard deviation of one experiment. Each symbol represents one technical replicate. Cells were gated on Via-probe^-^ CD4^+^ Vα2^+^ Vβ6^+^. (**b**) Fraction 17 from (a) was digested with trypsin in solution. The resulting peptides were analyzed by mass spectrometry. MS/MS-derived sequence data were used to search for matches against the *P. goldsteinii* proteome (refseq_P_goldsteinii_np1_20170705) and hits were validated using Scaffold. The table in (b) shows all hits identified on Scaffold using a 99% probability and a cutoff of at least 2 peptides/protein. (**c**) Representation of *P. goldsteinii* β-N-acetylhexosaminidase gene locus *(P. goldsteinii* CL02T12C30 supercont1.1, HMPREF1076).

**Extended Data Fig. 5:**
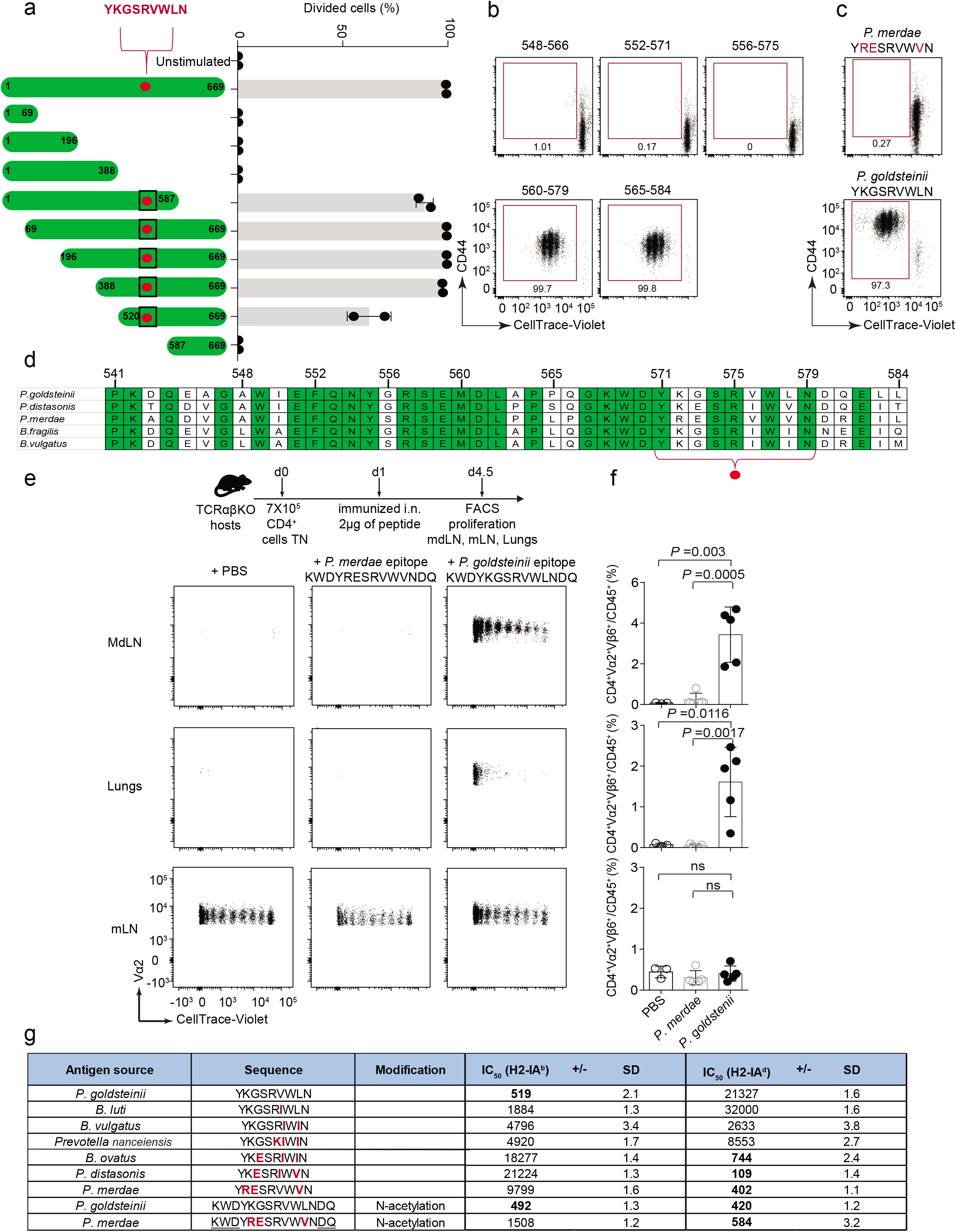
Truncations identify the segment of β-hexosaminidase recognized by the TN TCR. (**a**) Left: Truncations of β-N-acetylhexosaminidase (β-hex) were cloned and expressed in *E. coli.* The TN epitope is highlighted with a red dot. Right: Naïve CD4^+^ TN T cells were labeled with CellTrace-Violet and co-cultured with dendritic cells purified from B16-Flt3L– injected mice in the presence of the lysate derived from recombinant *E. coli* expressing the indicated β-hex truncations. 3.5 days later, dilution of CellTrace-Violet was assessed by flow cytometry. The graph shows the mean and standard deviation (SD) of one representative experiment of at least three independent experiments. Each symbol represents one technical replicate. Cells were gated on Via-probe^-^CD4^+^Vα2^+^Vβ6^+^. (**b**) Same as (a) but using synthetic peptides derived from *P. goldsteinii* β-hex as a source of antigen. The dot plots depict one representative replicate of three independent experiments. (**c**) Same as (b) but using minimal peptide epitopes derived from the *P. goldsteinii* and *P. merdae* β-hex. (**d**) Alignment of residues 541-584 of β-hex from different species (indicated on the left). Conserved positions are indicated in green. The TN epitope is highlighted with a red dot. (**e**) TCRαβKO hosts received 7X10^5^ naïve CD4^+^ TN T cells. The following day, the mice were immunized intranasally (i.n.) with 2 μg of peptide. 3.5 days later, the mediastinal lymph nodes (mdLNs), lungs and mesenteric lymph nodes (mLNs) were harvested and analyzed by flow cytometry. Dot plots show one representative mouse for each group. Cells were gated on CD45.2^+^CD4^+^Vα2^+^Vβ6^+^. (**f**) Graphs show the frequency (mean +/− SD) of TN cells (CD4^+^Vα2^+^Vβ6^+^) among CD45^+^ cells in the indicated organs of all mice analyzed in one experiment. Each symbol represents one mouse. (**g**) The table shows the IC_50_ and the geometric standard deviation (SD) for the binding of the indicated peptides to I-A^b^ and I-A^d^. IC_50_ values were assessed in at least 3 independent experiments. Bold values represent good binders (IC_50_<1000), while IC_50_ values between 1000-10,000 represent weaker binders. Peptide YRESRVWVN was flanked by sequences from the P*. goldsteinii* peptide (underlined in the table) to yield the longer *P. merdae* peptide KWDYRESRVWVNDQ. P values were calculated using unpaired one-sided Student’s t tests. P values <0.05 were considered statistically significant.

**Extended Data Fig. 6:**
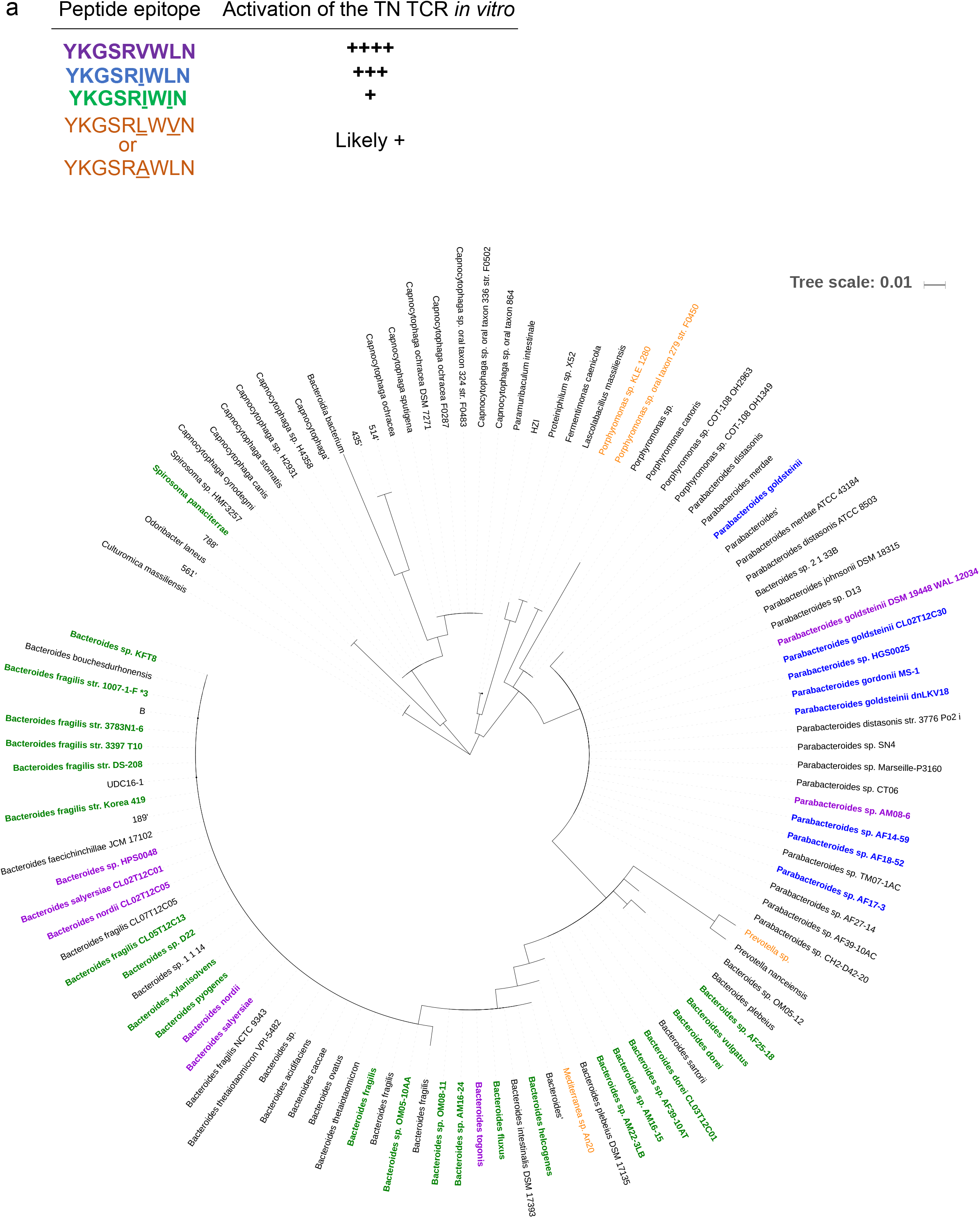
A broad range of *Bacteroidetes* species encode epitopes recognized by the TN TCR. (**a**) Phylogenetic tree constructed using the marker gene RNA polymerase β (RpoB) of species encoding a sequence similar to the TN epitope (YKGSRVWLN). Identification of species with similar epitopes was achieved by retrieving and aligning β-hexosaminidase (β-hex) sequences using Jackhmmer. Bold colored species encode the cognate sequence of TN T cells, orange species contain a likely activating sequence, while species in black do not. The bar shows the evolutionary distance that separates the bacterial species indicated (number of base substitutions per site).

**Extended Data Fig. 7:**
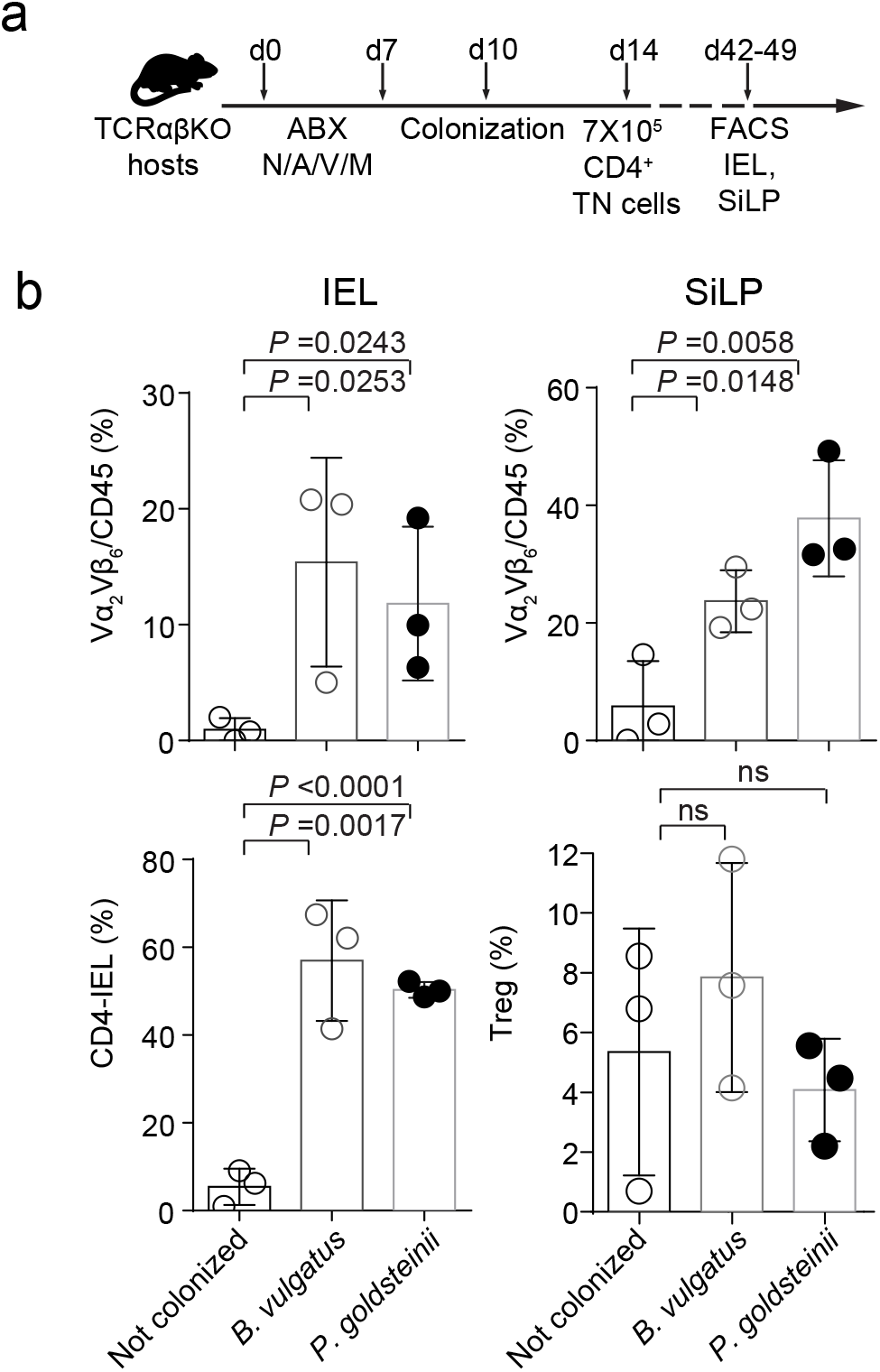
TN T cells differentiate into CD4_IELs_ upon colonization of immunodeficient hosts. (**a**) Experimental setup: TCRαβKO hosts were treated with antibiotics (ABX) for 7 days. On day 10, the host mice (n=3/group) were colonized or not with the indicated bacteria. On day 14, mice received 7X10^5^ naïve CD45.2^+^ CD4^+^ TN T cells. At day 42-49, the small intestine intraepithelial compartment (IEL) and the small intestine lamina propria (SiLP) were harvested and analyzed by flow cytometry. (**b**) Graphs shows all mice analyzed in one experiment. The frequency of CD4_IELs_ was assessed in the IEL and Tregs in the SiLP. Cells were gated on CD45.2^+^Aqua^-^CD4^+^ Vα2^+^Vβ6^+^TCRβ^+^TCRγδ^-^ in addition to CD8β^-^CD8α^+^Foxp3^-^ for CD4_IELs_ and CD8β^-^CD8α^-^Foxp3^+^ for Tregs. The graphs show the means +/− SD. Each symbol represents a single mouse. P values were calculated using unpaired one-sided Student’s t tests. P values <0.05 were considered statistically significant.

**Extended Data Fig. 8:**
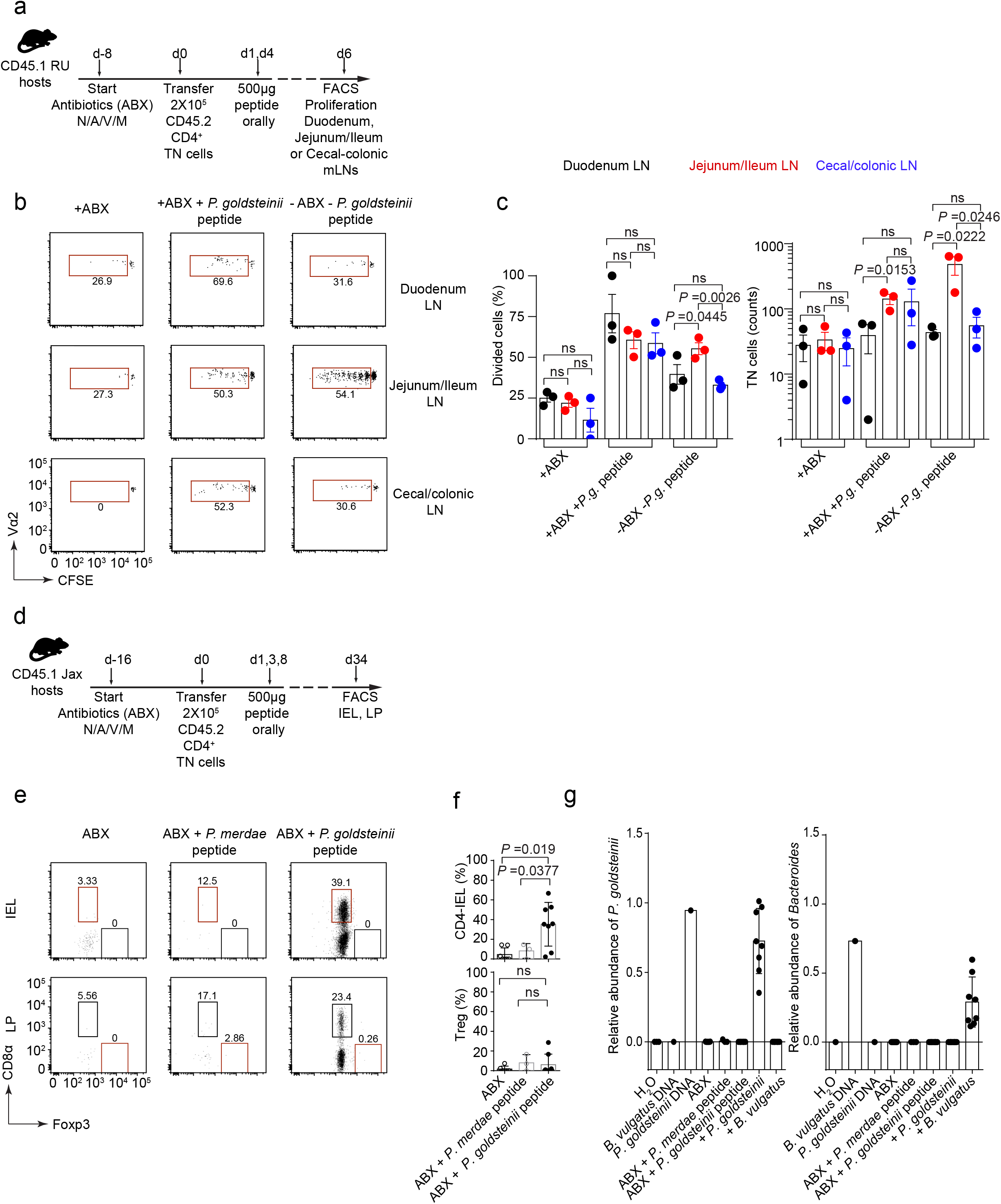
Oral delivery of cognate peptide is sufficient to induce proliferation and conversion of TN cells into CD4_IELs_. (**a**) Experimental setup: Congenically marked CD45.1 WT mice were treated with antibiotics (ABX) or not (n=5). Eight days later (d0), mice received 2X10^5^ CD45.1^+^CD45.2^+^ naïve CFSE-labeled CD4^+^ TN cells. At d1 and d4 post-transfer, some mice were given 500μg of β-hex peptide (KWDYKGSRVWLNDQ, n=5) by gavage. Six days posttransfer, T cell proliferation was assessed in the different mesenteric lymph nodes (mLNs). (**b**) Dot plots show CFSE dilution in live Aqua^-^CD4^+^Vα2^+^Vβ6^+^ CD45.1^+^CD45.2^+^T cells in the indicated mLN of one representative mouse. (**c**). Graphs show all mice analyzed in one experiment. (**d**) Experimental setup: Congenically marked CD45.1 WT mice were treated with antibiotics (ABX). Sixteen days later, mice received 2X10^5^ CD45.2^+^ naïve VioletTrace-labelled CD4^+^ TN cells. One, four and eight days post-transfer, the mice received 500μg of β-hex peptide from *P. goldsteinii* (KWDYKGSRVWLNDQ, n=8), from *P. merdae* (KWDYRESRVWVNDQ, n=3) or no peptide (n=7) by gavage. Thirty-four days post-transfer, expansion and conversion of TN cells into CD4_IELs_ (in the IEL) and Tregs (in the lamina propria, LP) was assessed by flow cytometry. (**e**) Dot plots show one representative mouse per group. Cells were gated on CD45.2^+^CD45.TAqua^-^ CD4^+^Vα2^+^Vβ6^+^CD8β^-^. (**f**) Graphs show all mice analyzed in two independent experiments shown in (e). (**d**) Colonization efficiency measured by qPCR. At the experimental end point (~4 weeks post-colonization), total fecal DNA was extracted from mice that received T cells, and colonized or not with the indicated bacteria (see e-f and Fig. 3l-n). qPCR was performed using *P. goldsteinii*-specific, *Bacteroides-specific* and 16s generic primers. The graphs show the relative abundance of *P. goldsteinii* and *Bacteroides* normalized to the total 16s (i.e Relative Abundance of *P. goldsteinii* = 2^-(CT *P. goldsteinii* -CT 16s)^) in all mice analyzed in two independent experiments. The experiments shown in Fig. 3l-n and Extended Data Fig. 8d-f were performed together and shown separately for clarity. The ABX group of mice represented in this figure is the same as the one shown in Fig. 3l-n. All mice were kept in microisolators with individual ventilation in the BCH facility. Each bar denotes the average+/− SD and each symbol represents a single mouse. The graphs show the means +/− SD and each symbol represents a single mouse. P values were calculated using unpaired one-sided Student’s t tests. P values <0.05 were considered statistically significant.

**Extended Data Fig. 9:**
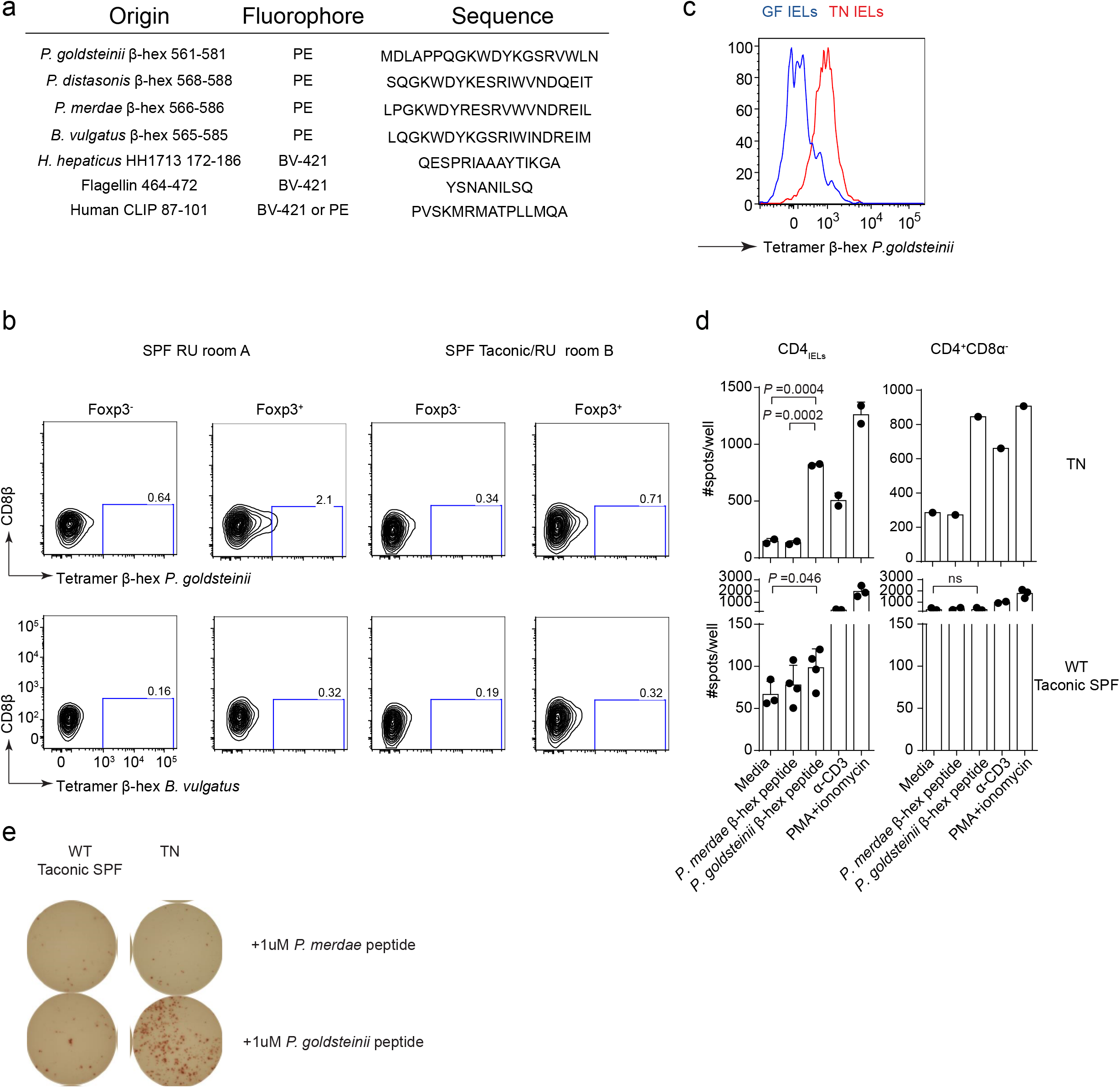
β-hexosaminidase MHCII tetramers identify antigen-specific CD4^+^ T cells in WT SPF mice. (**a**) List of tetramers used in this study. (**b**) Cells harvested from the lELs of WT germ-free (GF) or TN SPF mice were stained with MHCII *P. goldsteinii* β-hex tetramer. Cells were gated on CD45^+^CD4^+^Aqua^-^TCRβ^+^TCRγδ^-^. Histograms show one representative mouse of 3 different mice. (**c**) Cells from the mesenteric lymph nodes (mLN) of SPF mice housed at RU either bred at RU for many generations (SPF RU room A) or bought from Taconic 6 months prior to analysis and housed at RU in a different room (SPF Taconic/RU room B) were stained with the indicated tetramers. Cells were gated on CD45^+^CD4^+^Aqua^-^TCRβ^+^TCRγδ^-^ and Foxp3^+/−^ as indicated in the figure. (**d**) CD4_IELs_ from TN or Taconic SPF WT mice were harvested, sorted and co-cultured with splenic DCs isolated from TCRαβKO mice in the presence of 1 μM of the indicated peptides *(P. goldsteinii=KWDYKGSRVWLNDQ, P. merdae=* KWDYRESRVWVNDQ). After 18h of co-culture, IFNγ release was measured by ELISpot assay. Dot plots show one representative experiment for TN cells and all analyzed pools of WT mice (3-5 mice/experiment, mean +/− standard deviation). Cells were sorted on CD45^+^CD4^+^Aqua^-^TCRβ^+^TCRγδ^-^CD8β^-^ with CD8α^+^ or CD8α^-^ as indicated. (**e**) One representative ELISpot experiment as described in (d).

**Extended Data Fig. 10:**
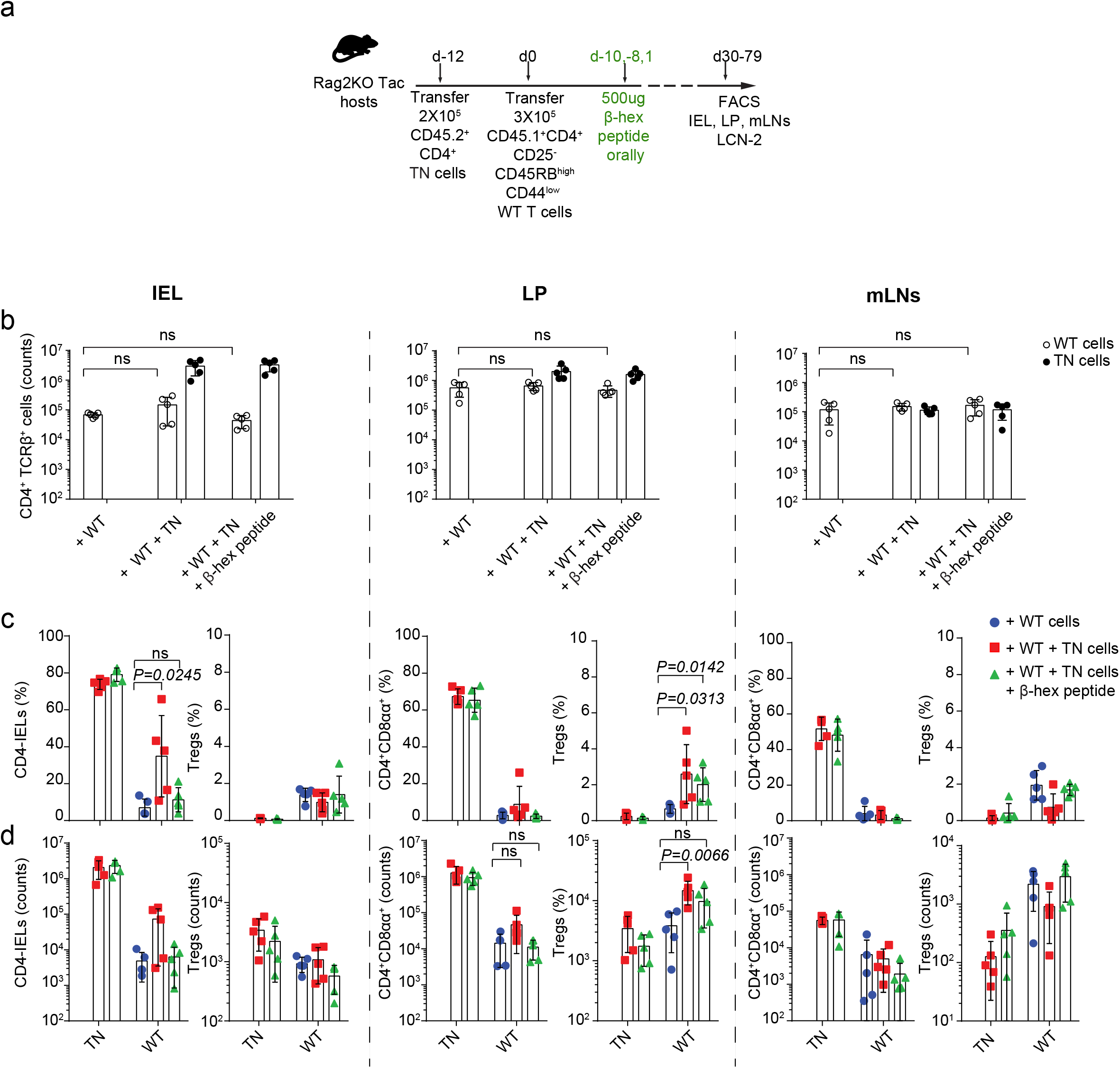
Migration and expansion TN and WT CD4^+^ T cells in lymphocyte-induced colitis. (**a**) Experimental setup: Taconic Rag2KO received 2X10^5^ naïve CD45.2^+^CD4^+^ TN cells or not. Twelve days later, the mice received 3X10^5^ naïve CD45.1^+^CD4^+^ WT cells (day 0). On days −10, −8 and +1 one group of mice received 500μg of *P. goldsteinii* β-hex peptide (KWDYKGSRVWLNDQ) orally by gavage. The mice were weighed every other day throughout the experiment. Mice were euthanized either when they lost 20% of their initial weight or at the end of the experiment. (**b**) Total number of TN (CD45.2^+^CD4^+^Aqua^-^TCRβ^+^) and WT cells (CD45.1^+^CD4^+^Aqua^-^TCRβ^+^) in the IEL (left), Lamina Propria (LP, middle) and mesenteric lymph nodes (mLN, right) in all mice analyzed in one experiment (n=5/group). (**c**) Frequency of CD4_IELs_ (CD4^+^Aqua^-^TCRβ^+^TCRγδ^-^CD8β^-^CD8α^+^Foxp3^-^, CD45.1^+^CD45.2^-^WT, CD45.1^-^CD45.2^+^=TN) or Tregs (CD4^+^Aqua^-^TCRβ^+^TCRγδ^-^CD8β^-^CD8α^-^Foxp3^+^, CD45.1^+^CD45.2-WT, CD45.1^-^ CD45.2^+^=TN) in the IEL (left), LP (middle) and mLN (right). (**d**) Total number of cells in the indicated organs, as described in (c). (b-d) The graphs show mean +/− standard deviation and each symbol one single mouse. P values were calculated using unpaired one-sided Student’s t tests. P values <0.05 were considered statistically significant.

## References

1 Hooper, L. V., Littman, D. R. & Macpherson, A. J. Interactions between the microbiota and the immune system. Science 336, 1268–1273, doi:10.1126/science.1223490 (2012).

2 Rooks, M. G. & Garrett, W. S. Gut microbiota, metabolites and host immunity. Nat Rev Immunol 16, 341–352, doi:10.1038/nri.2016.42 (2016).

3 Round, J. L. & Mazmanian, S. K. The gut microbiota shapes intestinal immune responses during health and disease. Nat Rev Immunol 9, 313–323, doi:10.1038/nri2515 (2009).

4 Atarashi, K. et al. Induction of colonic regulatory T cells by indigenous Clostridium species. Science 331, 337–341, doi:10.1126/science.1198469 (2011).

5 Sujino, T. et al. Tissue adaptation of regulatory and intraepithelial CD4+ T cells controls gut inflammation. Science 352, 1581–1586, doi:10.1126/science.aaf3892 (2016).

6 Bilate, A. M. et al. Tissue-specific emergence of regulatory and intraepithelial T cells from a clonal T cell precursor. Sci Immunol 1. eaaf7471, doi:10.1126/sciimmunol.aaf7471 (2016).

7 Tanoue, T. et al. A defined commensal consortium elicits CD8 T cells and anticancer immunity. Nature 565, 600–605, doi:10.1038/s41586-019-0878-z (2019).

8 Skelly, A. N., Sato, Y., Kearney, S. & Honda, K. Mining the microbiota for microbial and metabolite-based immunotherapies. Nat Rev Immunol 19, 305–323, doi:10.1038/s41577-019-0144-5 (2019).

9 Sefik, E. et al. MUCOSAL IMMUNOLOGY. Individual intestinal symbionts induce a distinct population of RORgamma(+) regulatory T cells. Science 349, 993–997, doi:10.1126/science.aaa9420 (2015).

10 Lathrop, S. K. et al. Peripheral education of the immune system by colonic commensal microbiota. Nature 478, 250–254, doi:nature10434 [pii] 10.1038/nature10434 (2011).

11 Yang, Y. et al. Focused specificity of intestinal TH17 cells towards commensal bacterial antigens. Nature 510, 152–156, doi:10.1038/nature13279 (2014).

12 Xu, M. et al. c-MAF-dependent regulatory T cells mediate immunological tolerance to a gut pathobiont. Nature 554, 373–377, doi:10.1038/nature25500 (2018).

13 Chai, J. N. et al. Helicobacter species are potent drivers of colonic T cell responses in homeostasis and inflammation. Sci Immunol 2, doi:10.1126/sciimmunol.aal5068 (2017).

14 Ivanov, II et al. Induction of intestinal Th17 cells by segmented filamentous bacteria. Cell 139, 485–498, doi:10.1016/j.cell.2009.09.033 (2009).

15 Linehan, J. L. et al. Non-classical Immunity Controls Microbiota Impact on Skin Immunity and Tissue Repair. Cell 172, 784–796 e718, doi:10.1016/j.cell.2017.12.033 (2018).

16 Bilate, A. M. & Lafaille, J. J. Induced CD4+Foxp3+ regulatory T cells in immune tolerance. Annu Rev Immunol 30, 733–758, doi:10.1146/annurev-immunol-020711-075043 (2012).

17 Shale, M., Schiering, C. & Powrie, F. CD4(+) T-cell subsets in intestinal inflammation. Immunol Rev 252, 164–182, doi:10.1111/imr.12039 (2013).

18 Reis, B. S., Rogoz, A., Costa-Pinto, F. A., Taniuchi, i. & Mucida, D. Mutual expression of the transcription factors Runx3 and ThPOK regulates intestinal CD4(+) T cell immunity. Nat Immunol 14, 271–280, doi:10.1038/ni.2518 (2013).

19 Cervantes-Barragan, L. et al. Lactobacillus reuteri induces gut intraepithelial CD4(+)CD8alphaalpha(+) T cells. Science 357, 806–810, doi:10.1126/science.aah5825 (2017).

20 Umesaki, Y., Setoyama, H., Matsumoto, S. & Okada, Y. Expansion of alpha beta T-cell receptor-bearing intestinal intraepithelial lymphocytes after microbial colonization in germ-free mice and its independence from thymus. Immunology 79, 32–37 (1993).

21 Langille, M. G. et al. Microbial shifts in the aging mouse gut. Microbiome 2, 50, doi:10.1186/s40168-014-0050-9 (2014).

22 Mucida, D. et al. Transcriptional reprogramming of mature CD4(+) helper T cells generates distinct MHC class Il-restricted cytotoxic T lymphocytes. Nat Immunol 14, 281–289, doi:10.1038/ni.2523 (2013).

23 Helgeland, L., Vaage, J. T., Rolstad, B., Midtvedt, T. & Brandtzaeg, P. Microbial colonization influences composition and T-cell receptor V beta repertoire of intraepithelial lymphocytes in rat intestine. Immunology 89, 494–501 (1996).

24 Wojciech, L. et al. Non-canonicaly recruited TCRalphabetaCD8alphaalpha lELs recognize microbial antigens. Sci Rep 8, 10848, doi:10.1038/s41598-018-29073-7 (2018).

25 Regnault, A., Cumano, A., Vassalli, P., Guy-Grand, D. & Kourilsky, P. Oligoclonal repertoire of the CD8 alpha alpha and the CD8 alpha beta TCR-alpha/beta murine intestinal intraepithelial T lymphocytes: evidence for the random emergence of T cells. J Exp Med 180, 1345–1358 (1994).

26 Regnault, A., Kourilsky, P. & Cumano, A. The TCR-beta chain repertoire of gutderived T lymphocytes. Semin Immunol 7, 307–319 (1995).

27 Regnault, A. et al. The expansion and selection of T cell receptor alpha beta intestinal intraepithelial T cell clones. Eur J Immunol 26, 914–921, doi:10.1002/eji.1830260429 (1996).

28 McDonald, B. D., Bunker, J. J., Ishizuka, I. E., Jabri, B. & Bendelac, A. Elevated T cell receptor signaling identifies a thymic precursor to the TCRalphabeta(+)CD4(-)CD8beta(-) intraepithelial lymphocyte lineage. Immunity 41, 219–229, doi:10.1016/j.immuni.2014.07.008 (2014).

29 Chapman, C. G. et al. Characterization of T-cell Receptor Repertoire in Inflamed Tissues of Patients with Crohn,s Disease Through Deep Sequencing. Inflamm Bowel Dis 22, 1275–1285, doi:10.1097/MIB.0000000000000752 (2016).

30 Dewhirst, F. E. et al. Phylogeny of the defined murine microbiota: altered Schaedler flora. Appl Environ Microbiol 65, 3287–3292 (1999).

31 Rakoff-Nahoum, S., Paglino, J., Eslami-Varzaneh, F., Edberg, S. & Medzhitov, R. Recognition of commensal microflora by toll-like receptors is required for intestinal homeostasis. Cell 118, 229–241, doi:10.1016/j.cell.2004.07.002 (2004).

32 Lagkouvardos, I. et al. The Mouse Intestinal Bacterial Collection (miBC) provides host-specific insight into cultured diversity and functional potential of the gut microbiota. Nat Microbiol 1, 16131, doi:10.1038/nmicrobiol.2016.131 (2016).

33 Grondin, J. M., Tamura, K., Dejean, G., Abbott, D. W. & Brumer, H. Polysaccharide Utilization Loci: Fueling Microbial Communities. J Bacteriol 199, doi:10.1128/JB.00860-16 (2017).

34 Liu, X. et al. Alternate interactions define the binding of peptides to the MHC molecule lA(b). Proc Natl Acad Sci U S A 99, 8820–8825, doi:10.1073/pnas.132272099 (2002).

35 Wexler, A. G. & Goodman, A. L. An insider,s perspective: Bacteroides as a window into the microbiome. Nat Microbiol 2, 17026, doi:10.1038/nmicrobiol.2017.26 (2017).

36 Eddy, S. R. Accelerated Profile HMM Searches. PLoS Comput Biol 7, e1002195, doi:10.1371/journal.pcbi.1002195 (2011).

37 Poyet, M. A library of human gut bacterial isolates paired with longitudinal multiomics data enables mechanistic microbiome research. In revision. (2019).

38 Esterhazy, D. et al. Compartmentalized gut lymph node drainage dictates adaptive immune responses. Nature 569, 126–130, doi:10.1038/s41586-019-1125-3 (2019).

39 Powrie, F., Leach, M. W., Mauze, S., Caddle, L. B. & Coffman, R. L. Phenotypically distinct subsets of CD4+ T cells induce or protect from chronic intestinal inflammation in C. B-17 scid mice. Int Immunol 5, 1461–1471 (1993).

40 Nutsch, K. et al. Rapid and Efficient Generation of Regulatory T Cells to Commensal Antigens in the Periphery. Cell Rep 17, 206–220, doi:10.1016/j.celrep.2016.08.092 (2016).

41 Geuking, M. B. et al. Intestinal bacterial colonization induces mutualistic regulatory T cell responses. Immunity 34, 794–806, doi:10.1016/j.immuni.2011.03.021 (2011).

42 Reis, B. S., Hoytema van Konijnenburg, D. P., Grivennikov, S. I. & Mucida, D. Transcription factor T-bet regulates intraepithelial lymphocyte functional maturation. Immunity 41, 244–256, doi:10.1016/j.immuni.2014.06.017 (2014).

## Material and method references

1 Bilate, A. M. et al. Tissue-specific emergence of regulatory and intraepithelial T cells from a clonal T cell precursor. Sci Immunol 1, eaaf7471, doi:10.1126/sciimmunol.aaf7471 (2016).

2 Rakoff-Nahoum, S., Paglino, J., Eslami-Varzaneh, F., Edberg, S. & Medzhitov, R. Recognition of commensal microflora by toll-like receptors is required for intestinal homeostasis. Cell 118, 229–241, doi:10.1016/j.cell.2004.07.002 (2004).

3 Valdez, Y. et al. Nramp1 drives an accelerated inflammatory response during Salmonella-induced colitis in mice. Cell Microbiol 11, 351–362, doi:10.1111/j.1462-5822.2008.01258.x (2009).

4 Maeda, H. et al. Quantitative real-time PCR using TaqMan and SYBR Green for Actinobacillus actinomycetemcomitans, Porphyromonas gingivalis, Prevotella intermedia, tetQ gene and total bacteria. FEMS Immunol Med Microbiol 39, 81–86 (2003).

5 Gomes-Neto, J. C. et al. A real-time PCR assay for accurate quantification of the individual members of the Altered Schaedler Flora microbiota in gnotobiotic mice. J Microbiol Methods 135, 52–62, doi:10.1016/j.mimet.2017.02.003 (2017).

6 Matsuki, T. et al. Development of 16S rRNA-gene-targeted group-specific primers for the detection and identification of predominant bacteria in human feces. Appl Environ Microbiol 68, 5445–5451 doi:10.1128/aem.68.11.5445-5451.2002 (2002).

7 Sidney, J. et al. Measurement of MHC/peptide interactions by gel filtration or monoclonal antibody capture. Curr Protoc Immunol Chapter 18, Unit 18 13, doi:10.1002/0471142735.im1803s100 (2013).

8 Cheng, Y. & Prusoff, W. H. Relationship between the inhibition constant (K1) and the concentration of inhibitor which causes 50 per cent inhibition (I50) of an enzymatic reaction. Biochem Pharmacol 22, 3099–3108 (1973).

9 Gulukota, K., Sidney, J., Sette, A. & DeLisi, C. Two complementary methods for predicting peptides binding major histocompatibility complex molecules. J Mol Biol 267, 1258–1267, doi:10.1006/jmbi.1997.0937 (1997).

10 Simon, M. D. et al. Rapid flow-based peptide synthesis. Chembiochem 15, 713–720, doi:10.1002/cbic.201300796 (2014).

11 Preheim, S. P., Perrotta, A. R., Martin-Platero, A. M., Gupta, A. & Alm, E. J. Distribution-based clustering: using ecology to refine the operational taxonomic unit. Appl Environ Microbiol 79, 6593–6603, doi:10.1128/AEM.00342-13 (2013).

12 Edgar, R. C. UPARSE: highly accurate OTU sequences from microbial amplicon reads. Nat Methods 10, 996–998, doi:10.1038/nmeth.2604 (2013).

13 Wang, Q., Garrity, G. M., Tiedje, J. M. & Cole, J. R. Naive Bayesian classifier for rapid assignment of rRNA sequences into the new bacterial taxonomy. Appl Environ Microbiol 73, 5261–5267, doi:10.1128/AEM.00062-07 (2007).

